# Representational geometry shapes noise-driven working memory errors

**DOI:** 10.64898/2026.11.23.768703

**Authors:** Matteo Alleman, Matthew Panichello, Timothy J. Buschman, W. Jeffrey Johnston

## Abstract

Humans and other animals have strictly limited working memory capacity. When asked to recall one or more presented items after a delay period, subjects will occasionally recall an apparently random item: a “guess” response. Where do these guesses come from? Using electrophysiological recordings from across monkey neocortex during the performance of a color working memory task, we show that the neural correlates of guess responses emerge during the memory delay – and that variable encoding precision makes guesses more likely, especially in one of our two animals. If guess responses emerge due to noise, we hypothesize that the detailed neural population geometry for color should shape the distribution of behavioral responses. We confirm this hypothesis by showing that the representational geometry specific to each animal is consistent with their idiosyncratic patterns of errors. Despite the often assumed robustness of neural population representations, noise appears to be a key bottleneck on the reliability of working memory.

## 1 Introduction

Working memory is limited; humans and other animals can remember only a handful of items reliably. For instance, when shopping for more than a few items in the grocery store, a shopper will typically refer to a list of the items because they cannot hold them all in working memory at once. These limitations on working memory lead to errors in behavior, such as the shopper forgetting to grab one of the items they were trying to remember. Decades of research has characterized working memory errors[1–12]. Small errors in memory recall are thought to arise from noise during memory encoding, maintenance, and report[9, 12, 13]. However, in challenging conditions, humans and other animals often produce apparently random – “guess” – responses. Our understanding of how these “guess” responses come about is incomplete. Previous work in psychology has posited that working memory may be limited in the number of items that can be simultaneously remembered[7, 14]. In this model, when asked to remember more items than the limit, some items will necessarily be forgotten or never encoded at all. If one of those non-encoded items is cued for recall, the participant makes a random guess. An alternative hypothesis posits that working memory is a continuous resource, typically some form of signal strength, that can be allocated across multiple items[1, 6] – either evenly or unevenly, with certain items being encoded with higher precision than others. If the item cued for recall was encoded with relatively low precision, the response could, in some cases, look like a random guess. Both of these theories have support from behavioral experiments[3, 14].

The neural substrate of working memory is believed to be the collective activity of large populations of neurons firing persistently[15, 16] (or sporadically[17–19]) to signal the remembered items. Electrophysiological work has shown that different items are typically represented in distinct subspaces of neural population activity[13, 20], consistent with an item limit. Other work shows that increasing the number of items to be remembered decreases the average information in single neurons about a specific item[21], consistent with the resource model. Continuous attractor dynamics in the neural population are thought to stabilize representations across the delay period. Relatively small noise would then drive diffusive dynamics on the attractor manifold[22–24], thus producing (small) working memory errors. As expected from these diffusion dynamics, prior experimental work in monkeys has shown that drift in the neural population representation across the delay period in a motor working memory task predicts small response deviations[12]. However, diffusive dynamics on a continuous attractor manifold cannot, by itself, explain guess responses[6–8, 13, 25].

Despite their extensive behavioral characterization, and the extensive neurophysiological study of working memory, it is still unknown precisely what failures of neural computation produce guess responses. In particular, the dynamics of target- and response-related information in neural population activity has yet to be characterized on trials with guess responses. Prior work has shown that there is decreased information about the target stimulus on trials with a missed change detection[21], which could make the representation of the target stimulus more vulnerable to corruption by noise or could be associated with eventual forgetting of the target stimulus entirely – both of which could lead to guess-like responses and are consistent with prior variable precision models of behavior[1, 3]. Recent theory and analysis of behavioral data has outlined an alternative: large noise during the memory delay could produce guess responses[8, 25–27]. In this theory, large noise drives both the diffusive dynamics that produce small, local errors and occasional large-magnitude “threshold” errors to non-local parts of the representational manifold[26]. In contrast to prior theories, this would mean that the geometry of this representational manifold links the size of these local errors to the frequency of non-local errors[26, 28, 29]. While models based on this theory fit behavioral data[8, 25], it is not yet known whether neural population representations have the geometric structure predicted by the theory.

This explanation of guess responses as noise-driven is also complicated by recent work showcasing the exquisite precision of neural population representations. Neural population representations in primary sensory areas have been shown to be far more precise than the corresponding behavioral judgments of the animal[30], indicating that information can be encoded with high precision in at least some cases (though other work has shown that encoding is weaker on trials when an error occurs[21]). Recent theoretical work has shown that reliable working memory is possible with only a handful of neurons[31]. In agreement with these findings, recent work on the neural basis of a specific type of working memory error (the “swap” error) shows that those errors do not arise from noise during the memory delay. Instead, they arise when color information is moved from one representational slot to another[10]. These results suggest that maintenance of information in working memory could be largely reliable, while there is a bottleneck on reliable encoding or manipulation of information in working memory.

Here, we adjudicate between these different possibilities and provide evidence that guess responses are driven by noise during the memory delay and shaped by representational geometry. In monkeys performing a two-item working memory task, we show that guess responses are associated with an early, reliable representation of the correct color that only later is corrupted to become a strong representation of the erroneously reported color. We also show that, on average, the target color is encoded more weakly on guess trials than on trials with correct responses, especially in one of our two animals. We develop a simple ring attractor model that links the level of variability in encoding precision to the scaling of error magnitude with memory delay length. We verify this link in our data. Next, we further develop and test the theory of noise-driven guess responses that are shaped by representational geometry. Consistent with the theory, we find that both animals have geometries that are consistent with their behavioral error distributions on our main task, as well as on a related task. We further show that the detailed representational geometry also predicts animal behavior. Overall, our results provide support for the idea that guess responses are driven by noise and shaped by representational geometry.

## 2 Results

To understand the neural basis of working memory errors, we trained two macaque monkeys to perform a visual working memory task with color stimuli. The task required the animals to remember two randomly and independently selected colors over a delay period before they were cued to select one of the colors based on its location (i.e., upper or lower). After a second memory delay, the selected item was reported via a saccade to the matching location on a color wheel (fig. 1a and see *Retrospective task* in *Methods* for more details). The color wheel was randomly rotated on each trial, so the animals could not plan their saccade in advance. The colors were selected from a set of 64 colors that were equally spaced on an isoluminant ring in HSV color space (fig. 1a, right). The two animals learned to perform the task well (fig. 1b), but both still made large report errors on a significant fraction of trials (fig. 1b). If these behavioral responses occur solely due to random, noise-driven diffusion on a ring attractor, then the deviations from the target color should have a wrapped normal distribution. We found that the wrapped normal distribution explains only part of the response distribution (fig. 1b, “local diffusion”), consistent with prior work in humans[1, 3, 6–8, 11, 25, 32, 33]. To explain the full distribution, we modeled it as a mixture between this wrapped normal component and a uniform component (fig. 1b, “guess rate”), which represents the probability of a “guess” response that is random with respect to the target color. We also included a “swap” component, capturing trials on which the animal reported the distractor rather than the target color[10]. Both animals were best fit by models with moderate local diffusion (fig. 1c, left; Monkey E: 0.89 rad^2^to 0.95 rad^2^; Monkey W: 0.69 rad^2^to 0.74 rad^2^) and “guess” response rates significantly greater than zero (fig. 1c, right; Monkey E: 0.18 to 0.24; Monkey W: 0.12 to 0.17).

**Figure 1:**
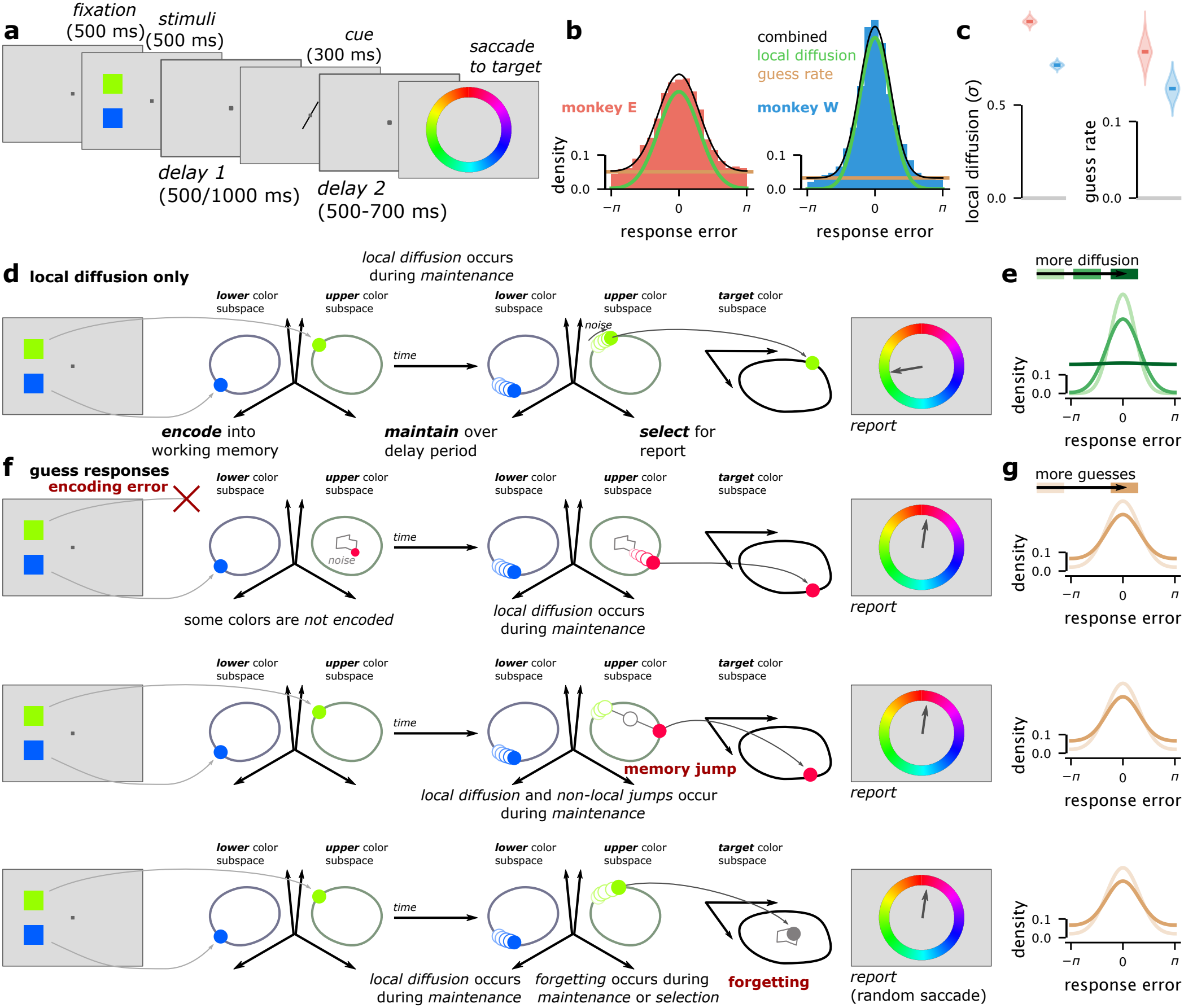
Working memory is limited and has multiple potential failure points. **a** Two animals performed a working memory task. The animals had to remember two randomly selected color stimuli across a delay period (delay 1). They were then cued to report one of the two colors based on its position (e.g., report the upper color; cue). There was a second delay period (delay 2) before the animal made a saccade to the appropriate color on a random rotated color wheel (saccade to target). **b** The distributions of responses for the two animals centered on the target color. **c** Following the convention in the field, we split the response distributions into two components: local errors, which are clustered near the target, and “guess” responses, which are uniformly distributed. **d** Schematic of the putative neural activity on a trial with a small local error (i.e., a “correct” trial). The schematic is based on prior work using these data[13]. **e** We focus on two possible sources of “guess” responses during this task. (top) A failure of encoding can produce “guess” responses, where only one (or neither) of the two colors is (are) initially encoded into working memory. Noise could still lead to the representation of a specific color, but that color would be random and lead to a random response. Local errors emerge due to noise during the delay period, but that noise is never large enough to produce a “guess” response. (middle) A failure of memory can produce “guess” responses, where noise during the memory delay jointly produces local errors and “guess” responses. Random, “guess” responses emerge naturally when the signal-to-noise ratio is low. (bottom) Guess responses may also arise from a forgetting process, where no color is represented in the target subspace because target color information is either never encoded (an encoding error, as above) or lost at some point during the memory delay. The animal’s guess responses would then be the result of a saccade to a random point on the color wheel.

Monkey E had both higher local diffusion (0.16 rad^2^to 0.23 rad^2^greater in monkey E than W) and a higher guess rate than monkey W (0.03 to 0.10 greater in monkey E than W). Unless otherwise stated, the ranges provided are 95 % confidence intervals for the mean.

While both animals performed this task (Monkey E: 13 sessions, Monkey W: 10), we recorded single and multi-unit neural activity simultaneously from posterior parietal cortex, frontal cortex, motor cortex, posterior inferotemporal cortex, and visual area 4 (posterior parietal cortex: 10 to 39 units; motor cortex: 18 to 53 units; prefrontal cortex: 22 to 57 units; V4 and posterior IT: 15 to 33 units; frontal eye fields: 2 to 23 units, where the ranges give the minimum and maximum number of simultaneously recorded neurons across sessions, and see *Electrophysiological recordings* in *Methods* for more details). We found a similar timecourse of trial information in frontal, parietal, and visual regions (fig. S1 and see *Region-level timecourse analysis* in *Supplement* for more detail). While there were some differences across regions in individual animals, the quantification within each region is noisy and the differences typically do not replicate across animals. As a result, we focused on the combined population.

### 2.1 Two competing explanations for large working memory errors

Here, we set out to understand the neural basis of local errors and guess responses. Prior work with this task in monkeys[13] has established the neural activity that unfolds on trials with small, local errors. On these trials, the two colors are initially represented in distinct subspaces corresponding to the position at which they were presented (fig. 1d, upper and lower subspaces, “encode into working memory”). So, on an example trial with a green color presented in the upper position and a blue color presented in the lower position (fig. 1a and d), the upper and lower subspaces would have neural activity representing green and blue, respectively, across stimulus presentation and the first delay period. Then, once the cue is presented, the target color is selected from either the upper or lower subspace and represented in a new, target subspace (fig. 1d, “select for report”). In our example trial, this means that the green color represented in the upper subspace will be rotated into this new target subspace. On correct trials, the blue color will not be rotated into the target subspace, though prior work has shown that a distinct kind of behavioral error does occur during this rotation – a “swap” error – which manifests as an erroneous rotation of the non-target color into the target subspace[10]. We excluded trials with these swap responses from our analyses. Once the second delay period ends, the animal must then map their color representation in the target subspace to a motor action – a saccade to the correct position on the randomly rotated color wheel (fig. 1d, “report”). Throughout this process, noise in the neural system is expected to produce local diffusion leading to non-zero response errors (fig. 1e). Prior work has shown that small errors are well-explained by this diffusive process[12], and that the population representation at the end of the memory period predicts these small deviations in the animal’s responses[12, 13].

However, errors in our task, as in many other working memory tasks[1, 3, 6–8, 11, 25, 32, 33], cannot be explained by diffusion alone (fig. 1c). How and when do the neural correlates of guess responses emerge? We work to distinguish between several competing hypotheses from the prior behavioral literature (fig. 1f)[7, 8]. One hypothesis is that guess responses emerge due to an encoding error, where one or both stimuli are either not encoded into working memory (fig. 1f, top, “encoding error”) or are encoded erroneously. Once this error is made at the time of encoding, it is reliably represented and transformed through the rest of the task (fig. 1f, top). This explanation would indicate that working memory itself is reliable, but that the encoding of items into working memory is unreliable or subject to significant constraints[7, 34, 35]. Prior electrophysiological work has shown support for this idea on a working memory change detection task[21], where the encoding of the target stimulus is weaker on trials where the change is missed.

An alternative hypothesis is that guess responses emerge despite correct encoding due to unreliable maintenance of information in working memory (fig. 1e, middle). In this case, blue and green are correctly represented in the upper and lower subspaces at first, but over the course of the first and second delay period, both representations undergo significant local diffusion and, in some cases, noise drives a jump to a non-local part of the representational manifold (fig. 1f, middle right, “memory jump”). These non-local jumps are sufficient to produce apparently random guess responses (fig. 1f, middle right). This hypothesis implies that neural noise is a significant factor in the reliability of some behaviors, which would be surprising given the fact that some sensory representations have been shown to be exquisitely reliable[30] and that working memory can, in principle, be noise robust[26] even with few neurons[36]. These two hypotheses are difficult to distinguish with behavioral data alone (fig. 1g), as they both lead to changes in the rate of guess responses. While both of these possibilities posit that the animal has a color in mind for report, a further possibility is that the animal completely forgets the target color on trials with guess responses and makes a saccade to a random point on the color wheel (fig. 1g, bottom). This could arise due to failures in either encoding or memory. One additional possibility is that the animal maintains a veridical representation of the target color through the trial until the report color wheel appears. Guess responses could then occur due to a failure of the animal to localize the target color on the report wheel. Our analyses will distinguish between all of these possibilities.

### 2.2 Behavioral errors emerge during the memory delay

To test these hypotheses, we began by separating the trials into correct (i.e., trials with responses that are likely to have come from the wrapped normal distribution centered on the target color; fig. 2a, green) and guess responses (i.e., trials with responses that are likely to come from a uniform distribution across all colors; fig. 2a, gold) using our behavioral mixture model (see *Behavioral mixture model* in *Methods* for details). This process excludes most trials with “swap” responses (i.e., trials with responses that are similar to the distractor color; fig. 2a, gray). We have outlined the neural dynamics on these swap trials in prior work[10].

**Figure 2:**
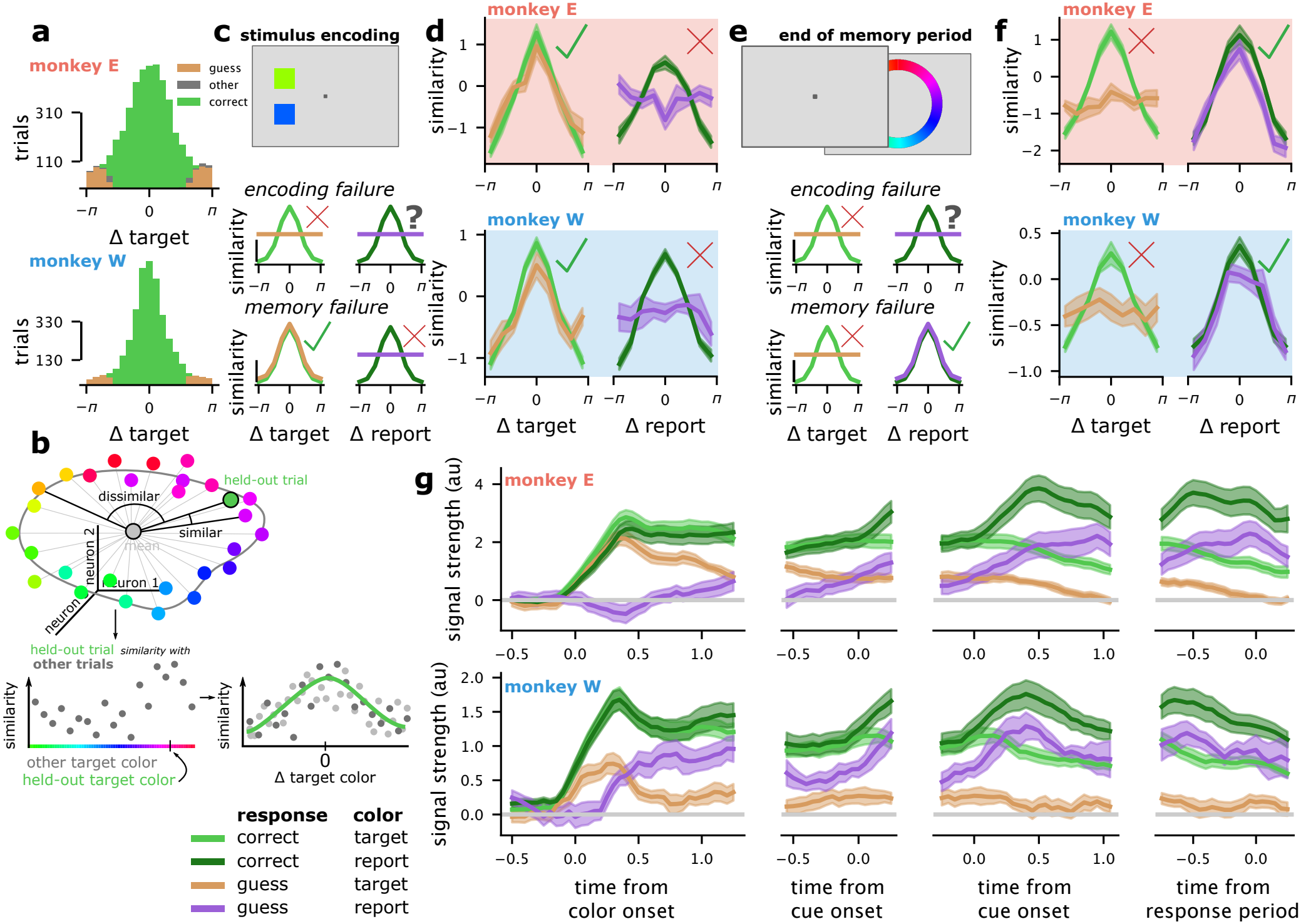
Single-trial error signatures primarily emerge during the memory delay period. a We split each trial into those with “correct” (i.e., small local error) and “guess” (i.e., random) responses, as well as a third category (“swap” errors) which have been investigated in prior work[10] and are excluded from this analysis. b We quantify color information in neural population activity (top) by taking the dot product similarity between a held-out trial and all other trials (bottom left). Then, we organize these pairwise comparisons by the difference in target color (bottom right) and average across held-out trials (green trace). **c** During stimulus encoding, an encoding failure would manifest as a strong representation of the target color on correct trials, but a non-existent or weak representation of the target color on guess trials (top left). The response color would also be represented on correct trials (on those trials, it matches the target), but it may or may not be represented on guess trials. In contrast, if encoding was reliable and errors happened later in the trial, then we expect strong representation of the target color on both correct and guess trials as well as no representation of the response color on guess trials. **d** The data is consistent with reliable encoding in both animals, where errors emerge later, as both animals have strong representations of the target color on both correct and guess trials as well as no representation of the response color on guess trials. **e** At the end of the memory period, an encoding failure would look the same as before, since the error had already occurred at the start of the trial. In contrast, a memory failure would now manifest as a strong representation of the target color on correct trials, but no representation of it on guess trials. In contrast, we expect to see a strong representation of the response color on guess trials. **f** The data is consistent with memory errors in both animals as the previously strong representation of the target color disappears on guess trials and is replaced by a strong representation of the response color. **g** The timecourse of color information across the three main periods of the task. This finer-grained analysis shows that target color information fades rapidly on guess trials, to be replaced by response color information well in advance of the response period. The target color information also never reaches the same maximum strength on guess as on correct trials, indicating that guess trials may already be more vulnerable to noise due to weaker color encoding at the start of the trial.

We developed a procedure for quantifying the similarity of the neural population representation at a given moment to a representation of the target color (fig. 2b; and see *Population representation similarity analysis* in *Methods* for more details). In brief, we take the neural population response from a single trial with a correct response (fig. 2b, top, outlined point). Then, we compute the dot product similarity between the activity on that trial and all other correct trials (fig. 2b, top and bottom left). A large, positive dot product between two trials means that the neural activity on those trials is similar (fig. 2b, top, “similar”) – i.e., pointing a similar direction in the high-dimensional population space – while a small or negative dot product means that the neural activity on those two trials is dissimilar (fig. 2b, top, “dissimilar”). We index each pairwise comparison between trials by the difference in their target colors (fig. 2b, bottom right) and repeat this procedure so that every correct trial is compared to all other correct trials (but never to itself). We bin these comparisons according to the difference between the target colors, then we take the average of the comparisons within each bin (fig. 2b, bottom right, green trace). This procedure quantifies how similar the neural representation is across correct trials with a given difference in target colors. If the target color is represented in population activity, then we expect similarity to be highest at the center of the function, where the target color across the compared trials is similar, and lowest at the sides, where the target color across the compared trials is maximally different.

We repeat this procedure for guess trials, by comparing the population responses on correct trials with the responses on guess trials. This allows us to leverage the larger number of correct trials to gain insight into the representations on guess trials, by making the assumption that the two kinds of trials share a common representation of color. If this assumption is violated, then we would find flat similarity functions for all analysis involving guess trials. We further tested whether guess trials are significantly different in other ways from correct trials by attempting to decode whether a trial will have a guess or correct response (fig. S2 and see *Decoding trial type* in *Supplement*). This decoder determined correct or guess response with significantly greater than chance performance only during the second delay period, prior to the response (monkey E: 0.55 to 0.57 decoding performance, *p <* 0.001, df = 12, monkey W: 0.53 to 0.55 decoding performance, *p* = 0.014, df = 9). This indicates that guess trials are only marginally different from correct trials, though it’s possible that other regions would provide a stronger generic error signal.

To distinguish whether guess responses occur due to failures of encoding or of memory, we analyzed the neural population activity during the stimulus encoding period of our working memory task (0 ms to 500 ms after stimulus onset, the entire display period of the stimuli; fig. 2c, top). If encoding failures explain guess responses, then we expect a strong encoding of the target color on correct trials (fig. 2c, middle left, green trace), but a weak or non-existent encoding of the target color on guess trials (fig. 2c, middle left, gold trace). Alternatively, if memory failures explain guess responses, then we expect a strong encoding of the target color on both correct and guess trials (fig. 2c, bottom left).

In both animals, we found strong representations of the target color on both correct and guess trials (fig. 2d, left). This finding is consistent with the memory failure hypothesis, since it indicates that information about the target color is initially present at stimulus encoding – and only later forgotten. However, in one animal, the representation of the target color was significantly weaker on guess trials than on correct trials (SEM range, monkey W: 0.71 to 1.11 au, *p* = 0.001, df = 9 weaker encoding of the target color on guess trials; the other animal trended in the same direction, monkey E: 0.18 to 0.64 au, *p* = 0.098, df = 12). This indicates that a weaker – rather than total failure of – stimulus encoding could still play a role in producing guess responses at least in one animal. We discuss the implications of this below.

Next, we analyzed neural population responses from the end of the second memory delay period (fig. 2e, top). The encoding failure hypothesis makes the same predictions as before: the target color should be represented on correct trials but not on guess trials. The memory failure hypothesis now makes the same prediction as the encoding failure hypothesis (fig. 2e, bottom left). If the target color is still represented at the end of the memory period on guess trials, the animal may make the error during the response period. Once the color wheel appears, the animal must both locate the target color on the randomly rotated color wheel and plan a saccade to that location. The retrieval, search, and motor planning processes could all, in principle, be error-prone and lead to guess responses. For instance, on some trials, the animal might fail to retrieve the target color from working memory or they might fail to rapidly locate it on the response wheel. Alternatively, our data simply may not capture neural population representations that are relevant for the behavioral report – e.g., the error may happen in some other brain region which we did not record from. In both animals, we found a strong representation of the target color on correct trials (monkey E: 2.42 to 3.08 au, *p <* 0.001, df = 12; monkey W: 0.84 to 1.18 au, *p <* 0.001, df = 9), but no representation of it on guess trials (fig. 2f, left; monkey E: 0.03 to 0.52 au, *p* = 0.283, df = 12; monkey W: *−*0.15 to 0.24 au, *p* = 0.810, df = 9). This indicates that the error in neural computation that produces guess responses happens prior to this report period.

Overall, our data provide support for the hypothesis that large working memory errors occur during the memory delay: The target color is initially represented on guess trials (fig. 2d, left), but that representation is lost by the end of the memory delay period (fig. 2f, left). Is any color represented on guess trials? On guess trials, the animal could forget the target color and make a random saccade that is either planned in advance or decided upon during the response period. Alternatively, if guess responses are driven by noise, the representation of the target color could be transformed across the delay period into a representation of a different color, which the animal then reports. To test for this second possibility, we used the same correlation analysis as above, but quantified similarity to the reported color rather than target color. While the reported color was not represented during stimulus encoding on trials with guess responses in either animal (fig. 2d, right; monkey E: *−*1.01 to *−*0.38 au, *p* = 0.048, df = 12, monkey W: *−*0.03 to 0.43 au, *p* = 0.410, df = 9), we found a strong representation of it at the end of the memory period (fig. 2f, right; monkey E: 2.14 to 3.02 au, *p <* 0.001, df = 12, monkey W: 0.71 to 1.00 au, *p <* 0.001, df = 9). This indicates that the animals do have a specific color in mind at the end of the memory delay – and that they do not make either a planned or random saccade during the response period. They reliably make a saccade to the color they have in mind, but it is the wrong color. This is consistent with the fact that, in these data, reaction times on correct trials are similar to reaction times on guess trials[10].

We then performed the same analysis in a sliding window across the entire time-course of a trial. Instead of showing the full average similarity of population representations to the target and response colors, we measure only the height of the function as an index of signal strength (fig. 2g, top). This analysis reveals different dynamics of error emergence across the two animals. In particular, the representation of the response color emerges later in monkey E than in monkey W (fig. 2g, bottom). This is consistent with the finding that the representation of the target color on guess trials is weaker relative to the representation of the target color on correct trials in monkey W, indicating that these representations may be more vulnerable to corruption by noise (explored in more detail below).

In addition, both animals appear to show an increase in the representation of the reported color around the time of the cue (fig. 2g, middle), though both the target color has started to be forgotten (SEM range for how much stronger the target color representation is on correct versus guess trials and one-sided, paired T-test: monkey E: 0.96 to 1.38 au, *p <* 0.001, df = 12; monkey W: 0.55 to 0.91 au, *p* = 0.001, df = 9) and, at least in one monkey, the reported color has started to be represented (SEM range for the strength of the representation of the reported color on guess trials, and one-sided T-test: monkey E: 0.21 to 0.77 au, *p* = 0.055, df = 12; monkey W: 0.34 to 0.61 au, *p* = 0.003, df = 9) prior to the appearance of the cue. This indicates that selection from working memory could amplify this error, similar to our prior work which showed that selection from working memory appears to be the moment that a different kind of working memory error (i.e., a swap error) occurs[10].

### 2.3 Errors are explained by neural noise and variable-precision encoding

To better understand the representational dynamics underlying errors in working memory, we turned to a highly simplified dynamical model of memory: a noisy ring attractor (fig. 3a). In this model, the representation of the one-dimensional, periodic color stimulus is embedded as a ring in a two dimensional space. From every point in this two-dimensional space, the dynamics of the system will move the representation toward the nearest point on the stable, unit-radius ring (fig. 3a, colored ring; see *Simplified ring attractor model* in *Methods*). Without noise, the representation would have a perfect memory: from any initial condition (except the exact origin) the representation would move to the nearest point on the ring and then remain there indefinitely. This model qualitatively replicates the dynamics of the ring attractor models used widely in neuroscience[31, 37–39], but in a reduced-dimensional setting that eases interpretation and visualization. The equations of the model are,

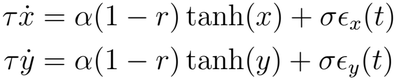

where *α* is a positive parameter controlling the strength of the attractive dynamics, *r* = *x*^2^ + *y*^2^is the current squared radius of the representation, *σ* is the standard deviation of the noise, and *ϵ*_*a*_(*t*) is Gaussian white noise, uncorrelated across *x* and *y*.

**Figure 3:**
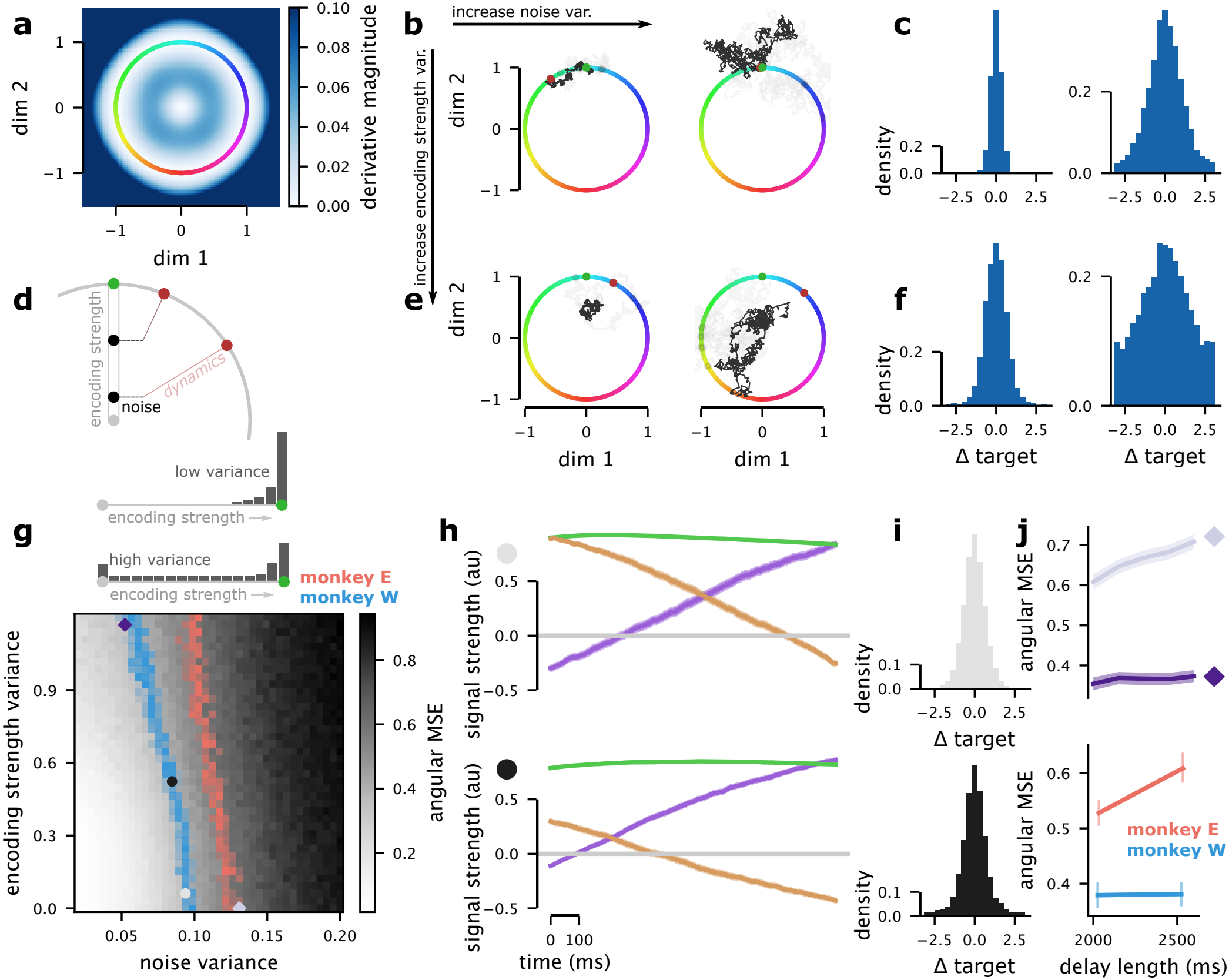
A simple attractor model explains guess-trial dynamics. **a** The dynamical landscape of our simple attractor model. The network has a stable ring of fixed points (colored ring) and a central, unstable fixed point (white center area). **b** Example simulations of this system with low (left) and high (right) noise. The target is always the green point, and the example trial reaches is closest to the red point at the end of a simulated delay period. **c** The error distributions for low (left) and high (right) noise. **d** The encoding strength shapes error the error distribution of the model. The same noise perturbation (dashed black line) causes a larger response error when the encoding strength is small (bottom) relative to large (top). **e** Example simulations of this system with low encoding strength and low (left) or high (right) noise. f The error distributions for low encoding strength and low (left) or high (right) noise. **g** (top) A schematic of the encoding strength distributions for low and high encoding strength variance. (bottom) The angular MSE of error distributions from models with different combinations of encoding strength variance and noise level. The colored lines show angular MSE values that match the angular MSE from the two animals. Note that each animal is has the same angular MSE as a variety of parameter combinations. **h** The dynamics of representational similarity from models with either low (top) or high (bottom) encoding strength variance. Higher encoding strength variance better replicates the representational similarity dynamics from monkey W (see fig. 2g, bottom left). **i** The error distributions from the models in h, and matched angular MSE values. **j** (top) The dependence of angular MSE on delay length for a model with high noise variance but low encoding strength variance (light purple, and see **g**) as well as a model with low noise variance and high encoding strength variance (dark purple, and see **g**). (bottom) The same as above but for the two animals.

We studied the memory performance of this model across a delay period. Without loss of generality, the model must always remember the state *x* = 0 and *y* = 1, or *θ* = 0. In our experimental setting, this is analogous to remembering a single, specific color. We then compared this target state, *θ*, to the actual _representation,_ ^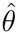^ after a delay period during which the memory was corrupted by noise and subject to the above dynamics. First, we visualized the single trial dynamics in both low and high noise (*σ*) conditions (fig. 3b). The representation clustered around the stable ring, but did not remain at a single point due to the random perturbations. With larger noise, the representation made larger excursions both away from and along the ring. The low-noise model had memory performance that is more precise than either of our animals (fig. 3c, left), while the high-noise model had qualitatively similar memory performance to our animals (fig. 3c, right).

The second parameter that we varied was the encoding strength of the remembered stimulus. In this setting, the initial condition of the model can be used to set the initial strength of the memory (fig. 3d). A weaker encoding will have lower signal-to-noise ratio in the early memory period, such that the same noise perturbation will produce a larger error if it happens early rather than late in the memory period (fig. 3d, bottom black point relative to bottom black point). For the same noise level, trials with a weaker encoding will therefore be more likely to have non-local memory errors than trials with a stronger encoding. This idea, often referred to in the human working memory literature as a variable precision encoding[1, 3], can be used to explain behavioral patterns of working memory errors. Here, we manipulated the encoding strength within the model on different trials (fig. 3d, high encoding strength; fig. 3e, low encoding strength; and similarly for fig. 3c, f) while studying how this changes the distribution of memory errors as well as the dynamics of representational similarity.

We set encoding strength to be between 0 (no encoding) and 1 (a full strength encoding, where the representation starts on the stable ring; fig. 3g, top). The encoding strength on a given trial was sampled from a truncated Gaussian distribution. The distribution has its peak likelihood at one (i.e., a full strength encoding). The variance then controls the distribution across lower encoding strengths (fig. 3g, top, “low variance” and “high variance”), where a higher variance means that lower encoding strengths are more likely. Then, we calculated the angular mean squared-error (MSE) from 5000 simulated trials from models with different noise and encoding strength variances (fig. 3g, bottom). This shows that angular MSE depends on both parameters. We also compared the angular MSE from these models with the angular MSE from our two animals (fig. 3g, red and blue shaded areas). The angular MSE from each animal is consistent with many different parameter choices. In particular, the same angular MSE can be achieved by decreasing the encoding strength variance as the noise variance increases.

We hypothesized that a difference in the variance of encoding precision can explain the weaker initial representation of the target color on guess trials in monkey W versus E (fig. 2f, left). This theory predicts that monkey W would have higher encoding strength variability than monkey E. Indeed, we find that the representational similarity dynamics in our models – computed in the same way as on the real neural data – qualitatively replicated the difference across the two animals (fig. 3h, top: low encoding strength variance, bottom: higher encoding strength variance), even while holding angular MSE constant (fig. 3i).

Then, we tested a second prediction of our model. For a constant angular MSE, higher encoding strength variability should mean that more large errors are made early rather than later in the trial. This is because a representation with low encoding strength is more vulnerable to noise than one with high encoding strength (fig. 3d), but the dynamics of the model will “forget” the initial encoding strength over time as it relaxes toward the stable ring. Our prior result indicates that monkey E, on average, has greater encoding strength than monkey W. It follows from this that the representation of the response color would emerge sooner in monkey W (or a lower encoding strength variability model; fig. 2f, top left; fig. 3h, top) than in monkey E (or a higher encoding strength variability model; fig. 2f, bottom left; fig. 3h, bottom).

In a task with variable delay lengths, this theory makes another prediction. When errors occur primarily due to noise during the memory delay rather than due to encoding strength variance, then the angular MSE should increase as the memory delay increases, consistent with noise-driven diffusion on an attractive manifold. However, if noise is relatively small but the encoding strength variability is high, then the early noise perturbations will dominate the ultimate angular MSE (fig. 3d). In this case, the angular MSE would not scale with small changes in delay length. We demonstrated these two regimes in our model, where the average angular MSE across different delay lengths is matched and delays range from 2000 ms to 2500 ms, as in our experiment (fig. 3j, top). These two regimes also matched the conditions of our two animals. Monkey E had small encoding strength variance but high noise (e.g., fig. 3h, top) while monkey W had large encoding strength variance but small noise (e.g., fig. 3h, bottom). Thus, the theory predicted that monkey E should show increased angular MSE with an increase in delay length, while monkey W should not. This is precisely what we observed (fig. 3j, bottom).

These modeling results both reinforce and elaborate upon the results of our analysis in the previous section. They show that differences in encoding strength can explain the differences that we observe in representation dynamics between the two animals: Monkey E has more consistent encoding strength, while monkey W has more variable encoding strength. Then, we use our simplified ring attractor model to show that the difference in encoding strength variability we observe neurally is sufficient to explain the difference in scaling of behavioral errors with memory delay that we observe between both animals. This indicates that the variable encoding precision is behaviorally relevant.

### 2.4 Neural population geometry could shape working memory errors

Our prior results provide evidence that large working memory errors occur due to noise and, to a de gree, how strongly the target is encoded. To further test this hypothesis, we now develop a stronger link between the structure of the neural population representation and the statistics of the behavioral responses. In particular, we hypothesized that the representational geometry for color in this task will constrain the distribution of behavioral responses. This builds on previous theoretical and behavioral work in humans[8], which showed that behavioral response distributions can be modeled as emerging from a particular neural population geometry[25]. Here, we tested if the actual neural population geometry corresponds to the one predicted by behavior under this theory.

A neural population representation of color (fig. 4a) can be summarized by its average geometry (fig. 4b), which describes the representational geometry only in terms of the difference between two colors. That is, the similarity function that describes the geometry is written as a function of the difference between two colors Δ = *x − y* rather than the two colors themselves: *k*(Δ) rather than *k*(*x, y*). This average similarity function can be computed by averaging over all color pairs with difference Δ in color space,

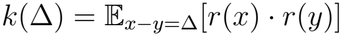

where *r*(*x*) and *r*(*y*) are the (noiseless) neural population representations of colors *x* and *y*, respectively.

**Figure 4:**
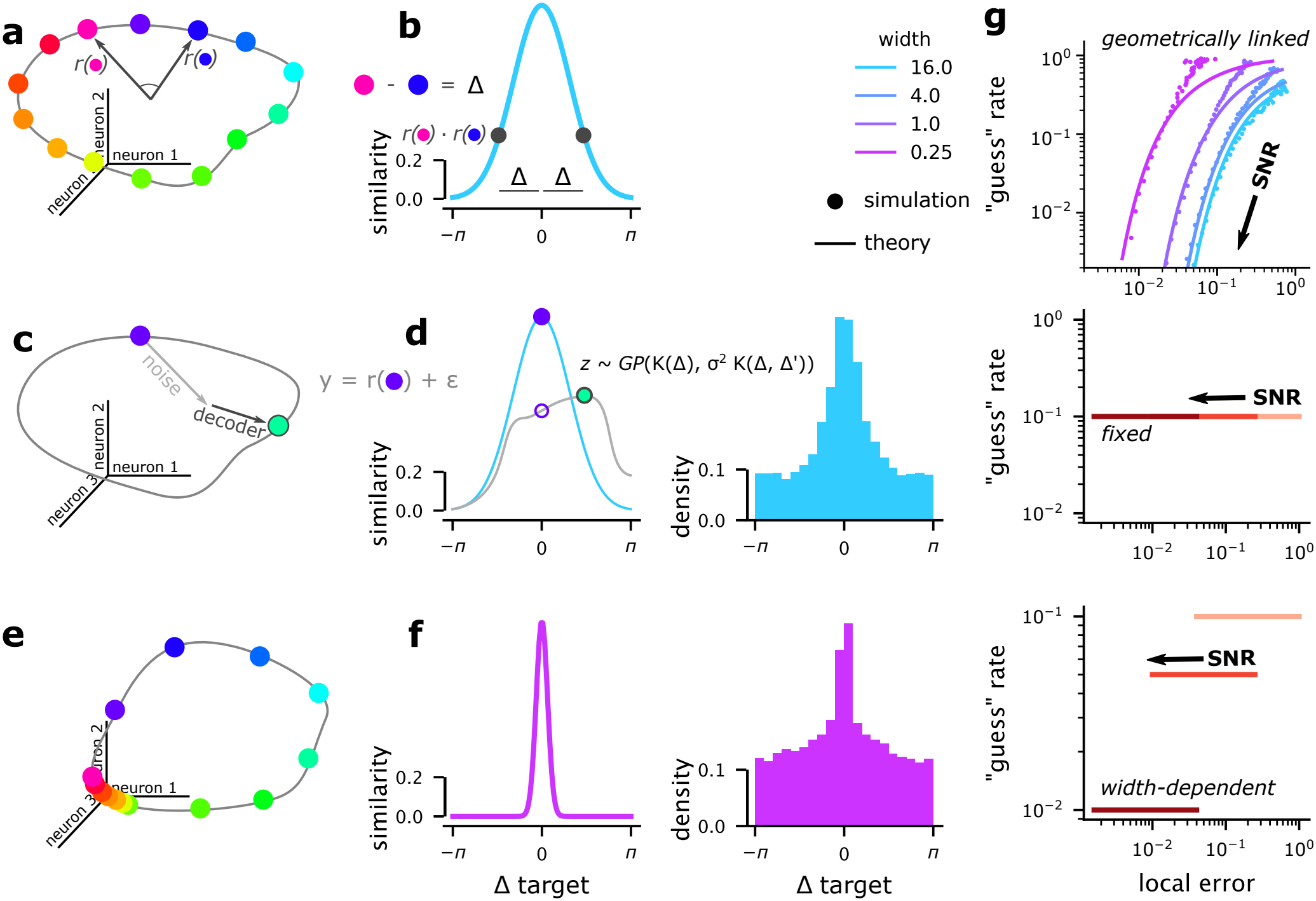
**A** theory establishing the correspondence between neural representational geometry and behavioral response distributions. a An example neural population representations of the color stimuli in the behavioral task. The population representation of different colors is given by *r*(.). **b** These population representations can be sumarized by an average similarity function, which describes how similar, on average, the population representations of two colors are, *r*(*x*) *· r*(*y*), given a particular distance from each other in color space, Δ = |*x − y*|. **c** Errors occur when noise corrupts a neural population representation (“noise”), and can lead to apparently random responses (“decoder”). **d** The effect of noise can be summarized in the similarity function, which now becomes a distribution of functions described by a Gaussian Process, *GP*(*K*(Δ), *σ*^2^*K*(Δ, Δ^*′*^)) (left). A draw from this process describes the similarity function on a given trial, for a given noise perturbation (gray line). The maximum of this trial function is the maximum likelihood response for the remembered color on that trial. Thus, the distribution of these maximums corresponds to a distribution of behavioral responses given that representational geometry and signal-to-noise ratio (right). **e** A schematic of a representational geometry that is higher-dimensional than the one in **a. f** A higher-dimensional representational geometry has a distinct similarity function (left) and, therefore, a distinct distribution of behavioral responses (right). **g** The geometric theory predicts that different representational geometries (colored lines) and SNRs will produce different combinations of local error magnitudes and “guess” response rates (top). Alternatively, the guess rate could be constant across different geometries and SNRs (middle) or depend on the geometry but not the SNR (bottom).

How would this neural population geometry influence behavioral errors? When the representation of a particular color, *r*(*x*), is corrupted by noise *r*(*x*) + *ϵ*, we can use a decoder to recover an estimate of the original color, 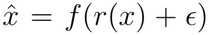 (fig. 4c). Consistent with prior work[25], we show that the color that would be decoded by an optimal maximum likelihood decoder is given by

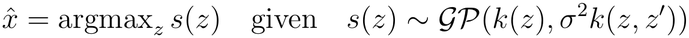

where *σ* is the standard deviation of the neural noise, which is assumed to be isotropic and Gaussian (fig. 4d, left; our theory can also be extended to Gaussian noise with any covariance structure). This equation shows that the estimate,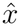, depends only on the average similarity function, *k*(Δ), and not on the full neural population representation, *r*(.).

In particular, each noisy representation corresponds to a sample from a Gaussian process, which has a mean and covariance structure determined by the average similarity function (fig. 4d, left, blue trace: similarity function and mean of the Gaussian process, gray trace: a sample from the Gaussian process corresponding to a single noisy color observation). The maximum of that sample then corresponds to the most likely original color (fig. 4d, left, cyan point). Many such samples produce a distribution of estimates that is qualitatively similar to the distribution of behavioral responses produced by our animals (fig. 4d, right). However, the shape of this distribution depends on the average similarity function. To illustrate this, we increased the dimensionality of the population representation in which the ring is embedded (fig. 4e), which leads to a narrower similarity function (fig. 4f, left) and a distinct distribution of behavioral responses (fig. 4f, right). This theory posits that the distribution of behavioral responses can be fully explained by the shape of the average similarity function, *k*(.), and the strength of the neural noise, *σ*.

We derive an improved analytical approximation for the error distribution given *k*(.) and *σ*. Compared to previous approximations in similar models [26], ours is better-suited to the high noise and broad similarity functions that we observe in the data. Importantly, the theory implies that the size of the local errors and the rate of guess responses will be linked by the average similarity function (fig. 4g, top, and see *Geometric theory of local error magnitude and guess rate* in *Methods* for details)[26, 28, 29]. This means that changing the average similarity function will change both the local error magnitude and guess rate and that the change in both will depend on the signal-to-noise ratio (fig. 4g, top). This need not be the case. If guess responses occur due to a fixed rate of encoding errors or forgetting, the guess rate would not depend on either the geometry or the SNR (fig. 4g, middle). Alternatively, the guess rate could depend only on the geometry, but not on the SNR (fig. 4g, bottom).

### 2.5 Neural population geometries are consistent with behavior across animals

The single and multi-units recorded as part of our experiments had heterogeneous color tuning (fig. 5a), making these individual units difficult to directly relate to behavior on their own. However, following from the theory above, we can abstract away single unit tuning by estimating the average similarity function of the neural population. We estimated the average similarity function by adapting an existing method for eigenfunction estimation (see *Kernel estimation* in *Methods* for details)[40]. Intuitively, we estimate the similarity function for a particular color difference Δ by taking the average dot product between all pairs of trials with target colors at that difference, Δ = *x − y* (fig. 5b) – note that this function is guaranteed to be symmetric around 0. We focused this analysis on the period just prior to behavioral report, and restricted it to trials where the animal made a likely correct response (p_correct_ *>* .5 and see fig. S7). As our prior results showed that the color mostly strongly represented at this time is the reported color rather than target color, we used this color to compute the geometry. This procedure produces a purely neural estimate of the average similarity function (fig. 5c, left, solid traces).

**Figure 5:**
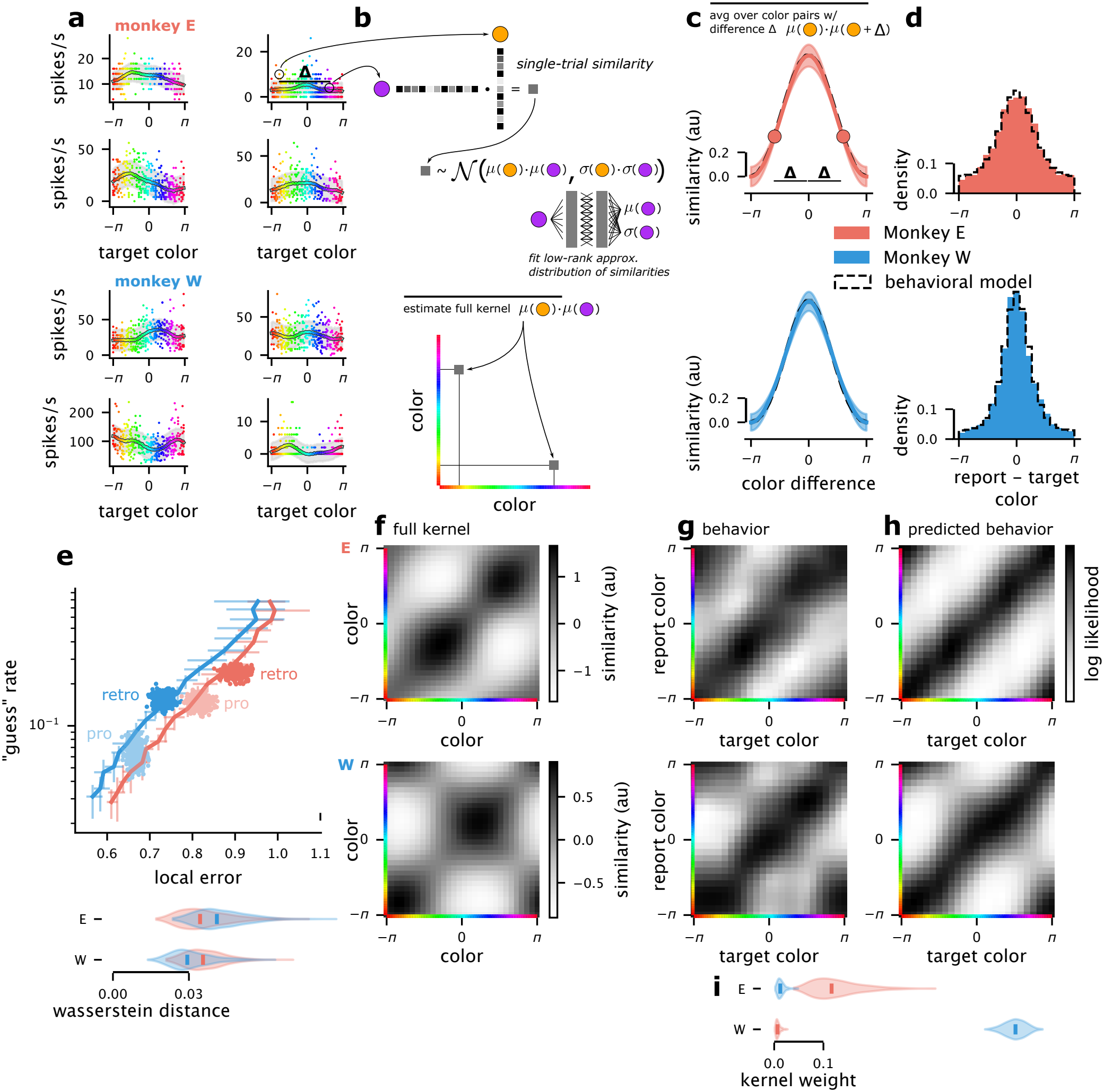
Neural response geometry shapes the behavioral response distribution. **a** Single unit examples from monkey E (top) and W (bottom). The individual points are responses from single trials. **b** The procedure for estimating both the average and detailed similarity functions. We use a neural network to estimate a low-rank approximation of the full distribution of similarities between trials as a function of their color representation. All population responses from single trials with small local errors are used. **c** The similarity function calculated from the neural population responses (solid line), compared with the similarity function inferred from the behavioral responses (dashed line). **d** The distribution of behavioral responses (solid histogram) and the distribution of behavioral responses predicted from the behavioral model (dashed line). **e** (top) The representational geometry from each animal (colored lines) define trajectories on the error plane defined in fig. 4g. These trajectories are consistent with the behavioral data on this task (point clouds). (bottom) Quantification of the fit between the geometry and behavioral responses. Low Wasserstein distance means a close fit between the distribution of responses predicted from the geometry and the true distribution of behavioral responses. **f** The detailed similarity function estimated from the population responses as in b for monkey E (top) and W (bottom). **g** The detailed distribution of behavioral responses for monkey E (top) and W (bottom). **h** The predicted responses from a behavioral model that weights the average and detailed similarity functions estimated from the neural data. **i** The weight given by the behavioral model to the animal’s detailed similarity function, contrasted with the weight given to the other animal’s detailed similarity function.

In parallel, we estimated the average similarity function that best explained the behavior of the animal. Here, we adapted a behavioral model introduced in prior work[8] to fit the behavioral response distribution using two parameters: a SNR and the width of the average similarity function (see *Behavioral similarity model* in *Methods* for details). The behavioral model follows the theory above, and finds the most likely SNR and function width given a set of behavioral responses. The behavioral responses sampled from the fit model matched the behavioral data (fig. 5c, right, solid: animal behavior, dashed black: model samples). This produced a purely behavioral estimate of the average similarity function (fig. 5c, left, dashed black trace).

We found that the purely neural and purely behavioral similarity functions match each other almost exactly (fig. 5c, left). The estimated tuning precision from the behavioral and neural data were not significantly different within animals (Monkey E: *−*0.17 rad^*−*1^to 0.08 rad^*−*1^ more behavioral than neural precision; Monkey W: *−*0.19 rad^*−*1^to 0.43 rad^*−*1^ more behavioral than neural precision). The behavioral tuning precision was significantly larger in monkey W than E (0.20 rad^*−*1^to 0.48 rad^*−*1^ more precision in Monkey W than E). The neural tuning precision was not significantly different across animals, but trended in the same direction (*−*0.23 rad^*−*1^to 0.43 rad^*−*1^ more precision in Monkey W than E).

Next, we took the average neural similarity function from each animal and simulated error distributions for a range of different signal-to-noise ratios levels (see *Noise-level simulations* in *Methods* for more details). This procedure produced a set of possible response distributions given a particular average similarity function, which we then decomposed into a local error magnitude and a “guess” rate (fig. 5e, blue and red traces) using the a behavioral mixture model. These traces are the geometric constraint implied by our theory (mirroring fig. 4g, top). The theory predicts that the local error magnitude and guess rate for both the task we have studied here as well as related tasks should lie on this line.

In our experiments, the animals performed the retrospectively cued task that we have already described (fig. 1a) as well as a prospectively cued task, in which the cue was presented prior to the color stimuli (see *Prospective task* in *Methods* for more details). Prior work with these data has shown that the prospective and retrospective tasks use aligned neural subspaces to represent color prior to the response period[10, 13]. Thus, assuming that this alignment indicates a shared representational geometry across tasks, the theory predicts that the average similarity function estimated in the retrospective task should define a constraint in local error magnitude-guess rate space across differing signal-to-noise ratios (i.e., a line, as in fig. 4g, top) that the prospective task obeys. This prediction need not be satisfied if guess responses arose as encoding errors and were therefore insensitive to the representational geometry.

For both animals, we estimated the average similarity function from the retrospective task and generate the constraint lines in local error magnitude-guess rate space following the theory (fig. 5e, lines). We then decomposed the behavioral response distributions on both the retrospective and prospective tasks into local error magnitude and guess response rate. This revealed that the behavioral data from both tasks tasks are consistent with the neural theory (fig. 5e). We then tested whether the geometry from one animal can provide a good fit to the behavioral data from the other animal. Here, we used Wasserstein distance to quantify the difference between the best fitting behavioral response distribution and the actual behavioral responses. While the behavior of each animal was slightly better explained by their own geometry rather than the other animal’s geometry (Monkey E’s geometry-constrained model fits monkey E’s behavior *−*0.02 to 0.04 more closely than monkey W constrained model, Monkey W’s geometry-constrained model fits monkey W’s behavior *−*0.02 to 0.03 more closely than monkey E constrained model), neither was significant.

While the average representational geometry provides a good explanation of behavior in both animals, we have not yet shown that differences in representational geometry lead to differences in behavioral response distributions. This is because both animals have similar average similarity functions. So, we move away from the average similarity function (which depends only on the difference between two colors) and to the full representational geometry, where different pairs of colors can have different similarities – even if they have the same distance from each other in color space. To estimate this geometry, we used the same novel method for low-rank kernel matrix approximation as before (see *Kernel estimation* in *Methods* for details). This gives us a reliable estimate of the detailed structure in the kernel matrix. For both animals, the estimated kernel is not strictly diagonal (fig. 5f) – and has distinct, block-like structure across different sets of colors. The detailed distribution of behavioral responses from each animal has some visual resemblance to this detailed kernel estimate (fig. 5g). Importantly, to estimate the detailed similarity function, we only use trials with small local errors (and a high posterior probability of being correct, see fig. S7 and fig. S8). Thus, this representational structure does not emerge due to behavioral sampling patterns.

To rigorously test whether the detailed structure of the kernel has an influence on behavior, we designed a behavioral model that explains behavior as a convex combination of the average similarity function from before and the color-specific full representational geometry (i.e., a horizontal slice of the kernel map shown in fig. 5f; see *Neural kernel behavioral model* in *Methods*). The degree to which the average and color-specific kernels are combined is controlled by a weight parameter. If this parameter is zero, only the average kernel is used. If the parameter is one, only the color-specific kernel is used. We fit this parameter to explain trial-by-trial behavioral responses, as before. The distribution of responses from this hybrid model (fig. 5h) fit the detailed behavioral response structure (fig. 5g). Further, for both animals, their own detailed representational geometry was significantly weighted by the model, while the other animal’s detailed geometry is not (fig. 5i). This means that an animal’s detailed representational geometry plays a significant role in shaping their behavioral responses – and does so in an animal-specific way (Monkey E’s detailed similarity function is weighted 0.07 to 0.27 more than monkey W’s function to explain monkey E’s behavior, Monkey W’s detailed similarity function is weighted 0.46 to 0.56 more than monkey E’s function to explain monkey W’s behavior). We replicated this result when resampling trials so that both animals have identical response distributions (fig. S8).

## 3 Discussion

Our work shows that guess responses in working memory are driven by noise and variable encoding precision as well as shaped by the representational geometry of color in the primate brain. Prior work has shown that a different kind of working memory error – the “swap” error – occurs during selection from working memory[10]. Other recent work has demonstrated that neural representation of sensory stimuli during encoding are extremely precise[30] as well as that ring attractors can be made reliable even with few neurons[31]. These three recent observations suggest a working memory system with a high degree of reliability in the maintenance of stimulus information, but which is highly constrained when manipulating that information. While our work shows that selection from working memory does appear to amplify the neural correlates of guess responses, the correlates exist prior to selection in both animals. Instead, our work provides evidence that working memory reliability is limited, in part, by significant noise in the neural system. This goes beyond prior work that has shown noise produces small, diffusive errors in other working memory tasks[12], and shows that noise can also lead to apparently random “guess” responses more frequently expected from noise-driven diffusion. We also show that the effect of this noise is amplified by variable precision encoding with predictable consequences for how error magnitude scales with memory delay length.

Given these noise-driven errors, we build on experimental and theoretical work that links the average representational geometry (that is, similarity between conditions as a function of their distance in stimulus space) to behavioral responses[8, 25]. First, our results support the hypothesis that the average similarity inferred from behavior corresponds to similarity in neural population geometry. Then, we go beyond this work by providing evidence that detailed representational geometry (that is, similarity in neural population representations of specific conditions) also plays a role in shaping behavioral responses. This result is particularly striking, since our data is distributed across several different regions in the monkey neocortex with only a few dozen units per region for every session. It is not obvious that such a dataset would be sufficient for estimating a behaviorally relevant, detailed (or even average) representational geometry. Our findings suggest that representational geometry is broadly consistent across the cortical hierarchy, and that widely distributed stimulus tuning and low-dimensional representations make it possible to estimate these behaviorally relevant geometries from relatively few neurons.

Prior experimental and theoretical work in decision-making tasks has demonstrated the importance of representational geometry for behavior[41–44]. This prior work focused on the qualitative affordances of a particular representational geometry: disentangled, or abstract representations enable generalization across contexts[41, 42, 45] while nonlinearly mixed representations enable the learning of complex task rules[43, 44]. While this work shows that representational geometry is disrupted on incorrect relative to correct trials[41, 42, 44], the nature of this disruption and the source of these errors was not determined. In contrast, our work shows that trials with correct and error responses have very similar geometry. Instead, noise drives erroneous representations within that geometry, while the geometry itself shapes which kinds of errors are likely to occur. Future work will be necessary to integrate these two different lines of work. Both, however, emphasize the importance of neural population geometry for structuring behavior.

Prior work in this same task paradigm has shown that the distribution of behavioral responses is can be explained by a memory system with a set of discrete attractors at different colors (as well as occasional, random guess responses)[46]. These discrete attractors warp the response distribution by making the cor responding, attractive colors more likely behavioral responses than than other colors. This phenomenon could be implemented through either dynamic or representational mechanisms. The representation of color could be perfectly uniform while the neural dynamics bias responses toward the attractor colors. Alternatively, the dynamics could be uniform, but the geometry of the neural representations could reflect these preferred colors. While the prior behavioral work showed some evidence for the former possibility by analyzing the structure of single trial biases[46], our work shows that the geometric structure necessary for the latter possibility is present. Thus, the brain may lie in between these two extremes, with both heterogeneous dynamics and representational geometry that work together to shape the distribution of working memory responses.

Our findings are in tension with longstanding theories of efficient coding and representational reliability. From a normative perspective, neural populations with narrow tuning for stimulus features have been shown to maximize the precision of decoding while minimizing the metabolic cost, in terms of spiking, of the neural code[26, 29, 47–49]. Recent work has extended this theory to neurons with heterogeneous response functions more similar to those we observe in our data (fig. 5a) with the same conclusion[28]. However, our data suggest neural responses are broadly tuned, which would only be optimal in the case of large noise. Consistent with the overall conclusions of our study, this suggests that neural representations in working memory are genuinely noisy and this noise leads to behavioral unreliability. Future work will be necessary to understand where this noise comes from and why behavioral capacity for working memory is so limited even though neural populations can, in theory, store information reliably.

Overall, our work provides strong evidence that unreliability in working memory behavior emerges due to noise in neural systems. We show evidence for this both in terms of the dynamics of representations on trials with correct and guess responses, as well as through detailed analysis of the representational geometry. Our results provide key insight into the bottlenecks on an essential behavioral capacity, and question established theory on the reliability of neural representations.

## Acknowledgements

We are grateful to Stefano Fusi, Ulises Pereira Obilinovic, Laura Driscoll, Ashok Litwin-Kumar, and other members of the Center for Theoretical Neuroscience at Columbia University and the Allen Institute for Neural Dynamics for very helpful discussions about earlier versions of this work. We are grateful to Kenji Muramoto, Drew Baughman, Arianna Pepin, and Allison Ong for administrative support. This work was supported by the following grants and foundations: NIH NINDS K99NS138578 (WJJ), Weill Neurohub Investigator Award (WJJ), Simons Foundation 542983SPI, Gatsby Charitable Foundation GAT3708, the Kavli Foundation, and the Swartz Foundation.

## Author contributions

MA and WJJ designed and implemented the analyses and theory. MP and TJB designed and executed the experiments. MA and WJJ made the figures and wrote the paper. MA, MP, TJB, and WJJ edited the paper.

## 4 Methods

### 4.1 Subjects

All experiments were performed in two adult Rhesus macaques (Macaca mulatta; 8-9 years old). Monkey E and W weighed 12.1 and 8.9 kg, respectively. The experimental procedures were approved by the Princeton University Institutional Animal Care and Use Committee and were in accordance with the policies and procedures of the National Institutes of Health.

### 4.2 Behavioral tasks

The behavioral tasks were implemented in Psychtoolbox and MATLAB (Mathworks). The visual elements were displayed on a Dell U2413 LCD monitor, which the animals viewed from a distance of 58 cm.

### 4.3 Retrospective task

The subjects initiated each trial by fixating a central point. Then, two randomly and independently selected color stimuli were displayed for 500 ms in fixed upper and lower positions. The color stimuli were chosen from a set of 64 colors, which were equally spaced along a photometrically isoluminant circle in CIELAB color space, centered at (*L* = 60, *a* = 6, *b* = 14) with a radius of 57 units. The two colors were sampled randomly. After the stimuli were presented, there was a 500 ms or 1000 ms delay period, followed by the presentation of an abstract cue for 300 ms. The cue indicates that either the upper or lower color is the target. We used two different cue sets: (1) lines oriented 45^*°*^clockwise or anti-clockwise from the horizontal meridian indicating the lower or upper position, respectively; and (2) a triangle or a circle, indicating the lower or upper position respectively. The data from both cue sets is combined for all analyses. After the cue, there was a 500 ms to 700 ms delay period. Finally, a randomly rotated, 2^*°*^-wide color wheel is presented, including all colors in the stimulus set. The animals must then make a saccade from their central fixation to the location on the wheel that matches the target color. The animals were rewarded according to the distance of their response (in degrees of visual angle) from the target color. The magnitude of reward was determined by a von Mises distribution centered at 0^*°*^ error with a standard deviation of 22^*°*^). They were not rewarded and received a short timeout (1 s to 2 s) if their response was greater than 40^*°*^ or 60^*°*^from the target, for Monkey E and W respectively. The animal had to make their response within 8 s, but the typical response time was within 200 ms to 300 ms.

### 4.4 Prospective task

The prospective task was the same as the retrospective task except that the presentation of the cue comes before the presentation of the color stimuli. After a 500 ms fixation period, the cue is shown for 300 ms. Then, there is a 200 ms to 600 ms delay period before the color stimuli (selected as before, from the same set of possible colors) are shown (500 ms). The stimuli are followed by a second delay period (Monkey E: 1000 ms to 2000 ms; Monkey W: 1300 ms to 2000 ms), before the same randomly rotated color wheel is shown for the response period.

The animals performed the retrospective and prospective tasks in blocks. The different cue sets for the retrospective task were defined as different blocks, for a total of three block types. Blocks transitions occurred after 60 correct trials on the prospective task and 30 correct trials on each cue set-specific retrospective task.

Throughout the experimental session, the eye position of both animals was tracked at 1 kHz using video-based eye-tracking (SR Research). Through the stimulus presentation, cue presentation, and delay periods the animal had to maintain their eye position within 2^*°*^ of a central fixation point. If they moved their eye outside of this window, the trial was aborted and the animal underwent a brief timeout.

Monkey E completed 11,131 trials over 13 sessions and Monkey W completed 9,865 trials over 10 sessions.

### 4.5 Electrophysiological recordings

The electrophysiological recordings are the same as performed in [10, 13]. In brief, the During the experiments, both animals were headfixed using a titanium headpost. The animals were each implanted with two titanium recording chambers, which were positioned to allow for recordings from LPFC, FEF, parietal cortex, and V4.

We used up to 80 simultaneously inserted expoxy-coated tungsten electrodes (FHB) to record single and multi-unit activity. After insertion, the electrodes were allowed to settle for 2-3 hours before recordings. Then, broadband (30 kHz) activity was recorded from each electrode (Blackrock Microsystems).

### 4.6 Behavioral models for working memory responses

We use a variety of models to explain the behavioral responses.

#### 4.6.1 Behavioral mixture model

We use the same hierarchical behavioral mixture model described in detail in [10], which builds on prior work[6, 7]. Briefly, we modeled the response error on each trial as a sample from a mixture model,

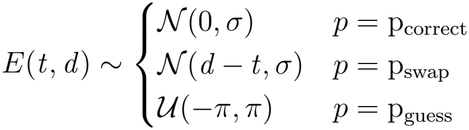

where the normal distributions are wrapped to be within *−π* and *π*. We fit this model in a hierarchical way, where *σ* and the other parameters for a single session are assumed to be drawn from a monkey specific distribution across sessions. This modeling approach provides us with a posterior probability that each trial is a correct, swap, or guess response. We then use this probability in our subsequent analysis, as described below and in the main text.

We initially fit a version of this model with wrapped Normal distributions in place of the Normal distribu tions shown above (fig. 1b). The wrapped Normal distribution describes the behavior of a representation that diffuses along a circular manifold under Brownian motion. For large diffusion noise (or a long in tegration time), the representation can make multiple trips around the ring. As *σ → ∞*, the wrapped Normal distribution converges to a uniform distribution on the ring. Since our data is best described by a relatively small *σ*, the fit from the wrapped Normal and from the Normal distribution are almost identical (fig. 1b). Due to this and for simplicity, we use the standard Normal distribution to compute posterior probabilities for classifying trials as correct or guess trials.

#### 4.6.2 Behavioral similarity model

We adapt the models described in [8, 11, 25] as alternate models for the behavioral responses. In contrast to the above mixture model, these models all assume that the behavioral response distribution emerges from a similarity function (often parameterized by its width, corresponding to how quickly the similarity between two different colors decays as a function of their difference) and the signal strength (equivalent to the magnitude of the similarity function or the inverse of the strength of the noise). These two parameters can produce response distributions that closely match the response distributions produced by the mixture model and, importantly, that provide a roughly equivalent match to behavioral data from humans performing working memory recall tasks[8, 11, 25].

In particular, given a similarity function *f*(Δ_*t*_;*w*), which is a function of the difference between the response color and target color and is parameterized by its width, *w*, the response error, Δ under the model is given by,

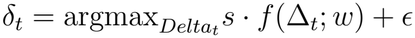

where *s* is the signal strength and *ϵ ~ N* (0, 1) is noise.

We modify this model to account for the probability of swap errors, as described above and treated in depth in [10], as a mixture component governed by p_swap_ – where the response is *δ*_*t*_ with probability 1 *−* p_swap_ and *δ*_*d*_ otherwise, with *δ*_*d*_ defined analogously to *δ*_*t*_.

The width *w*, signal strength *s*, and swap probability p_swap_ are all fit jointly using Pyro with stochastic variational inference.

#### 4.6.3 Neural kernel behavioral model

Since we are able to directly estimate the potentially relevant similarity function from neural data, we also fit a version of the similarity model above with a fixed similarity function – thereby reducing the number of parameters to two: signal strength *s* and swap probability p_swap_. We fit both an average similarity version of the model, where we use the same similarity function (the average given in fig. 5c) for all colors, as well as a full similarity model where the similarity function on each trial is given by *kf*(*c−c*_*t*_)+ (1*−k*)*g*(*c, c*_*t*_) with the kernel weight *k ∈* [0, 1] controlling the weight given to the color-specific similarity function *g* relative to the average similarity function *f*.

### 4.7 Population representation similarity analysis

We need a method for quantifying the color that is being represented in neural population activity at a particular time on a particular trial (the reference trial). Here, we develop a similarity-based measure. In particular, we take the dot product similarity (after z-scoring) between the population activity on the reference trial and all other trials. This gives us a collection of *T −* 1 similarities, where *T* is the number of trials.

To ask whether the neural activity is, on average, similar between trials with similar target colors, we can organize this collection of similarities by the difference between the target color on the reference trial and the target color on the compared trial – and then repeat this procedure where each individual trial is the reference trial, for a total of *T*(*T −* 1) similarities. We then average these similarities according to the difference between target colors. Further, to ask if this differs between correct and guess trials (e.g., fig. 2c), we can choose the set of reference trials to be only correct trials or only guess trials while the compared-to set of trials is all correct trials.

In the manuscript, we follow this latter procedure. For each session, we take all trials with a likely correct response (p_correct_ *>* .3 in fig. 2) and use these as reference trials. Then, we take the dot product similarity between each reference trial and all trials of interest (either the other correct trials or trials with likely guess responses), while keeping track of the difference in target (or response) color between the two trials. Note that the similarity of a trial is never taken with itself. We then bin these neural similarities according to the color difference (for instance, in fig. 2e we use 9 equally-sized bins across the interval [*−π, π*]), and average these bins across the different sessions. We perform this analysis on the z-scored neural responses.

This analysis is complicated by the fact that we expect the representation of the target color to differ during the first delay and stimulus presentation periods of the task, since the target trial will be in upper spatial location on half the trials and in the lower spatial location on the other half. Prior work has shown that these upper and lower subspaces are orthogonal to each other in neural population space[13]. So, we perform this analysis conditioned on cue. That is, we perform the analysis once for all trials with an upper cue (target color is in the upper position) and a second time for all trials with a lower cue (target color is in the lower position), then we average the results. We use this procedure in fig. 2b and throughout fig. 2f for consistency. In fig. 2e, we do not condition on cue. Conditioning on cue, or not, yields similar results after cue presentation.

### 4.8 Simplified ring attractor model

As stated in the text, we use a simplified ring attractor model with the following equations,

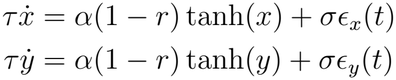

where *τ* is the time constant, *α* is a positive parameter controlling the strength of the attractive dynamics, *r* = *x*^2^ + *y*^2^is the current squared radius of the representation, *σ* sets the magnitude of the noise, and *ϵ*_*a*_(*t*) *~ N* (0, 1) is the noise.

From the equations, it is clear that *r* = 1 forms a stable ring manifold. The only other fixed point of the system is *x* = *y* = 0, which is unstable. We vary *σ* (the noise variance) and the initial encoding strength of the color memory in the ring system (i.e., the distribution of *r*_init_ *∈* [0, 1]. For the simulations we show in the manuscript, the attractive dynamics are chosen to be weak (relative to most choices of the noise magnitude), *α* = .01. Larger *alpha* would reduce the likelihood of guess errors in this model but would leave the diffusive (or local) errors at similar magnitude. The time constant is chosen to be moderate, *τ* = 5 ms; the other parameters could be rescaled to compensate for other values of *τ*.

We integrate these equations with Euler integration at 1 ms step size.

### 4.9 Geometric theory of local error magnitude and guess rate

Prior work[7] has assumed that the correct response precision and guess response frequency are indepen dent parameters. Recent work has observed that, if guess responses emerge due to noise on population representations, this would not be the case[8, 25, 33]. Instead, both correct response precision and guess response rate would depend on the underlying representational geometry and noise level.

This relationship has been shown analytically[26, 28, 29] but primarily outside of the working memory literature. Furthermore, those prior analyses have typically been done in the regime of relatively high SNR and narrow similarity functions; when those assumptions are violated, the approximations no longer capture the effects of large errors. So, here we make some corrections to account for these effects, and provide an alternative heuristic for the guess probability which works better for wide kernels.

Firstly, in the high-noise regime the response distribution has substantial mass across the entire [*−π, π*] range. As such, in this regime a theory whose predicted error is unbounded will begin to fail. To correct for this, we replace the standard Cramer-Rao bound on the MSE, *MSE ≥ F I*(*k, σ*)^*−*1^, with an _analogous bound on the mean cosine error (MCE) given by 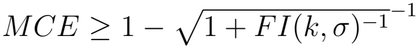_ which is more appropriate for periodic variables. Similarly, the union bound that [26] use to estimate the threshold error probability can exceed 1 when the noise variance is large, and so we derive another heuristic which always outputs a probability.

Secondly, wide similarity functions blur the distinction between local and guess errors, since the ‘local’ component covers nearly the entire circle. As such, we connect the geometry to the mixture model parameters via an intermediate concept of ‘threshold’ errors. More detail on this, and of the improved heuristics, can be found in S5.

#### 4.9.1 Noise-level simulations

To validate the theory developed in the previous section, and for our comparison to behavioral data, we simulate the response distributions for a variety of SNRs at different lengthscale widths (including the one estimated from the neural data).

We generate a population of neurons with Gaussian process response functions with lengthscale matched to the lengthscale estimated from the neural population. For a given SNR, we use the theory above to define a Gaussian process over likelihood functions on individual trials, then sample 4000 trials from that likelihood function and convert the likelihoods to discrete responses (by taking the argmax). Then, given this distribution of responses, we use a mixture model to estimate the correct response precision and guess rate. We average the correct precision and guess rate for a given SNR across 20 different sampled populations.

The mixture model is a simplified form of the hierarchical mixture model described above. We do not simulate separate sessions, so it is no longer hierarchical. Because we are simulating the response errors for a single stimulus, there are only correct and guess responses, swap errors cannot occur in this setting. The model is written and fit in Pyro using stochastic variational inference. The fitting is iterated for 5000 steps, the learning rate is 0.01. The model is fit with an independent Normal guide, which neglects any correlations between the two parameters.

### 4.10 Kernel estimation

We estimate the full (and average) kernel using the same core procedure, then average it to produce the reduced form of the kernel that depends only on color difference (e.g., in fig. 5c).

In general, for *T* trials, we have a *T × T* kernel matrix *K*, where each element *K*_*ij*_ is the dot product between the z-scored neural activity on trials *i* and *j*. We hypothesize that there is a *D*-dimensional, noisy function of color *r*_*i*_ *~ N* (*f*(*θ*), Σ(*θ*))) with *r*_*θ*_ *∈ R*^*D*^, where Σ(*θ*) is a scalar function of color – which implicitly makes the covariance matrix we estimate diagonal. This reflects the assumption that trial-to-trial noise is independent and does not depend on color proximity. Given this function and noise structure, we can sample *r*_*i*_ corresponding to the *θ* observed in our experiment that will then reproduce *K*. That is, *f*(*θ*) is a latent representation of color that we estimate directly from the kernel matrix itself – or, directly from the geometry of color representations rather than from the individual neural responses.

This problem is a less-constrained version of eigenfunction estimation, since we need not require the *D* dimensions of *f*(.) to be orthogonal or unit. To fit *f*(.) we adapt an eigenfunction estimation procedure known as NeuralEF [40], which uses an artificial neural network to approximate the function. However, while our problem is less constrained, it is also significantly noisier: We do not expect that, when the same color is shown on two different trials, both trials will have same response. Thus, we introduce the probabilistic machinery from above, and fit the mean and standard deviation of the function variationally using Pyro, by maximizing the likelihood of the data samples under the function distribution.

To determine the appropriate number of dimensions, *D*, for the function, we use a cross-validated halving grid search procedure. In brief, we estimate functions for all of *D ∈* 1, 2, …, 50 dimensions in 5-fold cross validation using a relative small number of training epochs given, then take the best performing half of candidate functions (in terms of average log-likelihood on the held-out data for each cross-validation fold). Then, we use the same procedure to estimate that set of functions again with an increased number of training epochs. The starting number of training epochs is 30 while the ending number is 150. We iterate this procedure until a single function estimate remains. This procedure is implemented in scikit-learn[50].

We use this procedure to estimate a single function for each experimental session, then average those estimates across sessions to arrive at the final kernel estimations shown in the main text.

## S1 Supplement

### S1 Region-level timecourse analysis

We investigate whether broad groups of regions appear to play different roles in the emergence of the neural correlates of guess responses. We replicate the timecourse analysis shown in the main text (fig. 2g) for subsets of single and multi-units from each of three broad region groups: frontal (including PFC and FEF), parietal (7ab), and visual (V4 and PIT). This analysis shows similar dynamics across each of the three region groups. The most striking difference is also the least surprising: The visual regions more strongly represent the colors during their presentation and the first delay period; while the parietal and frontal regions more strongly represent color in the second delay period.

Despite having a weaker representation of color during the second delay period, visual regions still show a loss of target color and gain of reported color information on guess trials – as seen in the combined population. Overall, the different region groups have a similar timecourse of target and reported color information on correct and guess trials as in the combined population. The differences that are apparent do not typically replicate across the two animals, complicating interpretation. Thus, we focus on the combined population.

### S2 Decoding trial type

To understand whether guess trials qualitatively differ from correct trials, we attempt to decode whether a particular trial results in a guess or correct response. To do this, we take the same data used in fig. 2, which is binned around the major task events and smoothed with a 500 ms boxcar window. Then, we label each trial as to whether it is likely to have a correct (p_correct_ *>* .5) or guess (p_guess_ *>* .5) response and train a SVM decoder with a radial basis function kernel to decode this label at each time point. The decoder is cross-validated using a shuffle and split procedure with 10 % of trials left out on every fold. The decoder and cross-validation procedure are implemented in scikit-learn[50].

The decoder reaches above-chance performance preceding the response period, after the cue is shown (fig. S2, top and bottom, right plots).

#### S3 Guess frequency across time

To determine whether the guess frequency changes significantly across time, we smooth the probability of a guess response with a 20 trial boxcar filter (fig. S3, left). We also plot the histogram of inter-guess intervals for p_guess_ *>* .4. Then, we fit this distribution with an exponential distribution and find good agreement (fig. S3, right). Together, these plots indicate that guesses occur with statistics similar to that of a Poisson process with constant rate across the session.

#### S4 Response color and previous trial responses

We ask whether the previous trial’s color influences the response on the current trial, and whether this influence is changed if the current trial is a guess response. For instance, one hypothesis would be that the guess response may simply replicate the previous trial’s response.

To do this, we first plot the two responses against each other in a scatter plot (fig. S6, red and blue points). Monkey E has a significant (but subtle) correlation between the prior and current response (Pearson *r* = .044 with *p* = .002). Then, we plot the same points but only when the current trial is a guess response (fig. S6, golden points). In this case, neither animal has a significant correlation between prior and current (guess) response.

**Figure S1:**
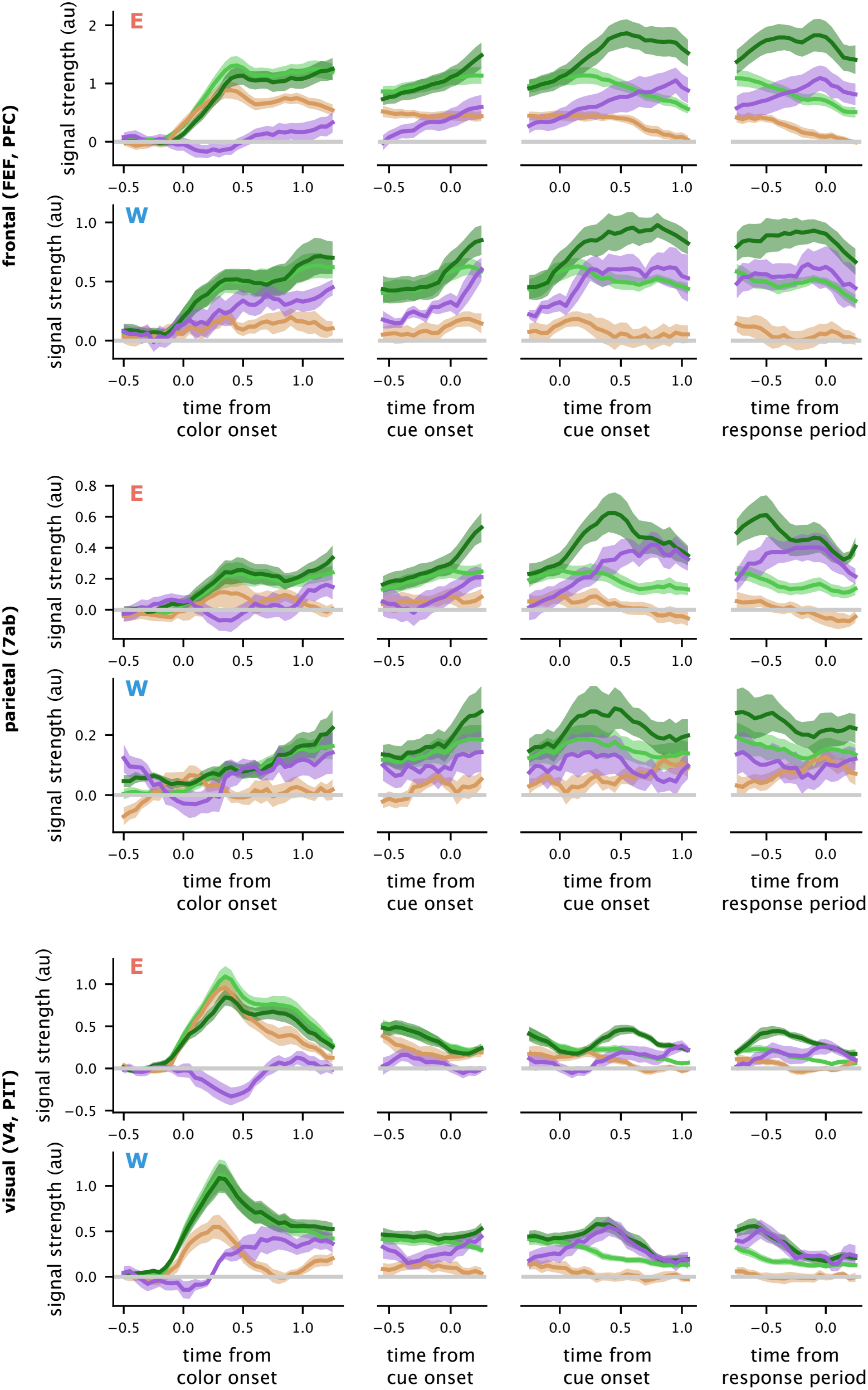
Analysis of the timecourse of target and reported color information on trials with correct and guess responses for multi-units and single neurons recorded from frontal (top), parietal (middle), and visual (bottom) regions. The analysis is the same as described in fig. 2g, but restricted to units from the indicated regions.

### S5 Details on model approximation

Here, we will explain the approximations used to compute mixture model parameters from the geometry and noise magnitude. Consider a periodic variable, *x*, which has a noiseless, high-dimensional neural representation, f(*x*). As explained in the main text, we model behaviour as maximum likelihood decoding from a working memory representation, f(*x*^*\**^)+*ε*, corrupted by gaussian noise *ϵ ~ N*(0; *σ*I). The noiseless representation f is assumed to have a similarity function, *k*(*x, x*^*′*^) = f(*x*) *·* f(*x*^*′*^), which is translation invariant, meaning *k*(*x, x*^*′*^) = *k*(*x − x*^*′*^). That assumption simplifies the maximum likelihood to

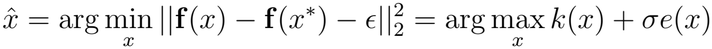

for a noise function *e*(*x*) *~ GP*(0, *k*(*·, ·*)) which is distributed as a Gaussian process with mean and kernel function given by *k*.

For simplicity, we will assume a specific kernel function which is unit-norm, *k*(0) = 1, and centered, 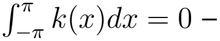 we use the modified von-Mises function 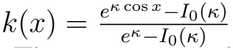. As *κ →* 0, this becomes a cosine kernel, cos *x*, which we call the “linear” regime. The same general approach could work for other kernels as well. Even with these assumptions, finding the distribution, or just its moments, of the maximiser of a GP-distributed function is not generally possible outside special cases. For instance, in the linear regime, the distribution of 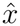 is known as the projected normal distribution. But otherwise, approximations are needed.

A familiar approach to this kind of maximum likelihood decoding is to make use of the Cramer-Rao lower bound on the variance of 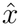, which is the inverse of the Fisher information (FI). However, this bound does not apply when 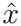 is periodic, since its variance cannot exceed 4*π*^2^ while the inverse FI can grow unbounded. Instead, we can use an analogous upper bound [51] on the first trigonometric moment of 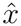, given by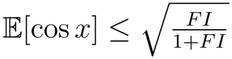. This is then a lower bound on the mean cosine error (MCE), 1 *−* E[cos *x*].

But the bound on error is not generally tight, and will severely underestimate the total variance of an estimator due to large errors. That is the problem that [26] address by estimating the probability of large, non-local errors that they call ‘threshold errors’. Briefly, the authors model a set of neurons with Gaussian tuning curves of width *w* and their preferred stimulus value, *x*_*i*_, uniformly spaced along the circle. They group these neurons into *n*_*ef f*_ = *π/w* effective groups (i.e. degrees of freedom) with mostly non-overlapping tuning. But the Gaussian kernel doesn’t respect the periodicity of *x*, and for wide nearly-linear von Mises kernels the concentration parameter, *κ*, is not an angular value that directly lets us count the effective number of independent groups. So, we chose the point 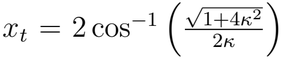, where the kernel turns from convex to concave, as a natural notion of width which generalises to wide kernel functions. The result is 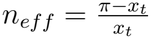 effective non-local groups.

**Figure S2:**
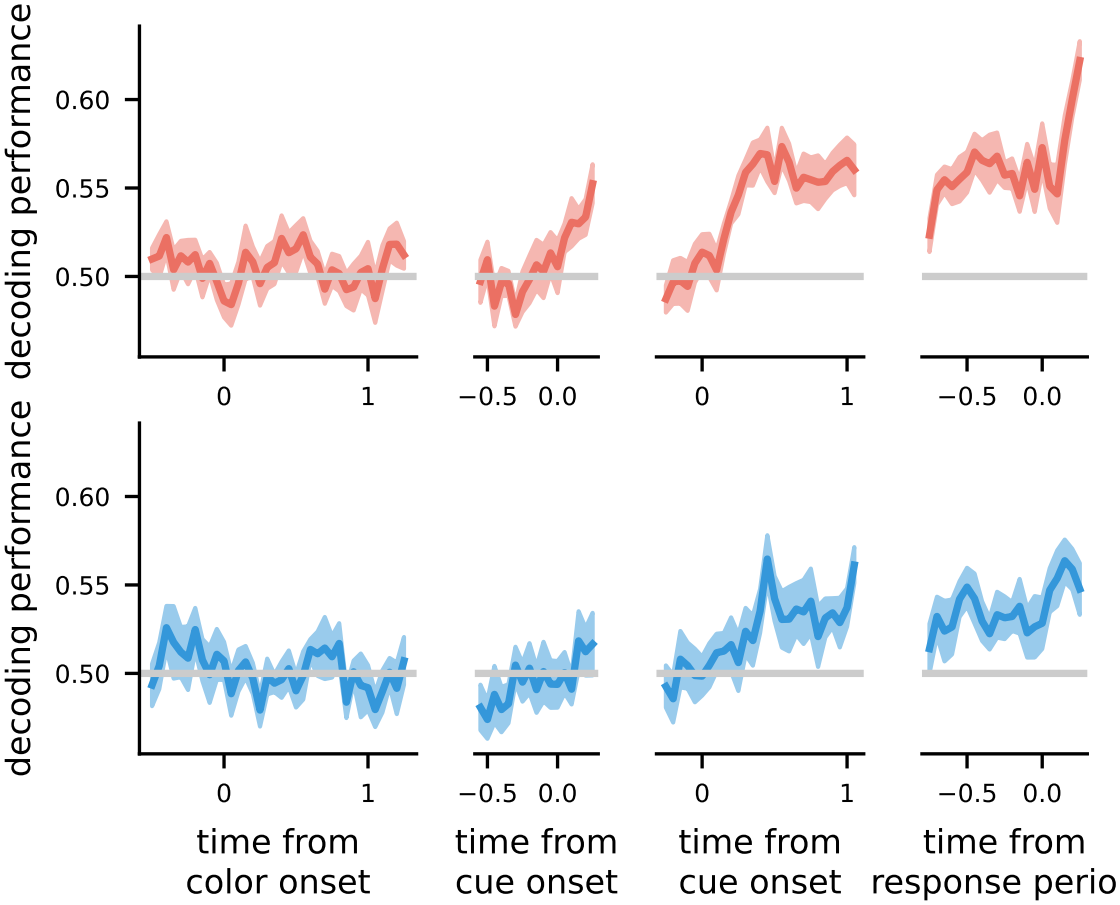
Decoding the response-type. The decoding performance for correct or guess behavioral response across the trial for monkey E (top) and W (bottom).

**Figure S3:**
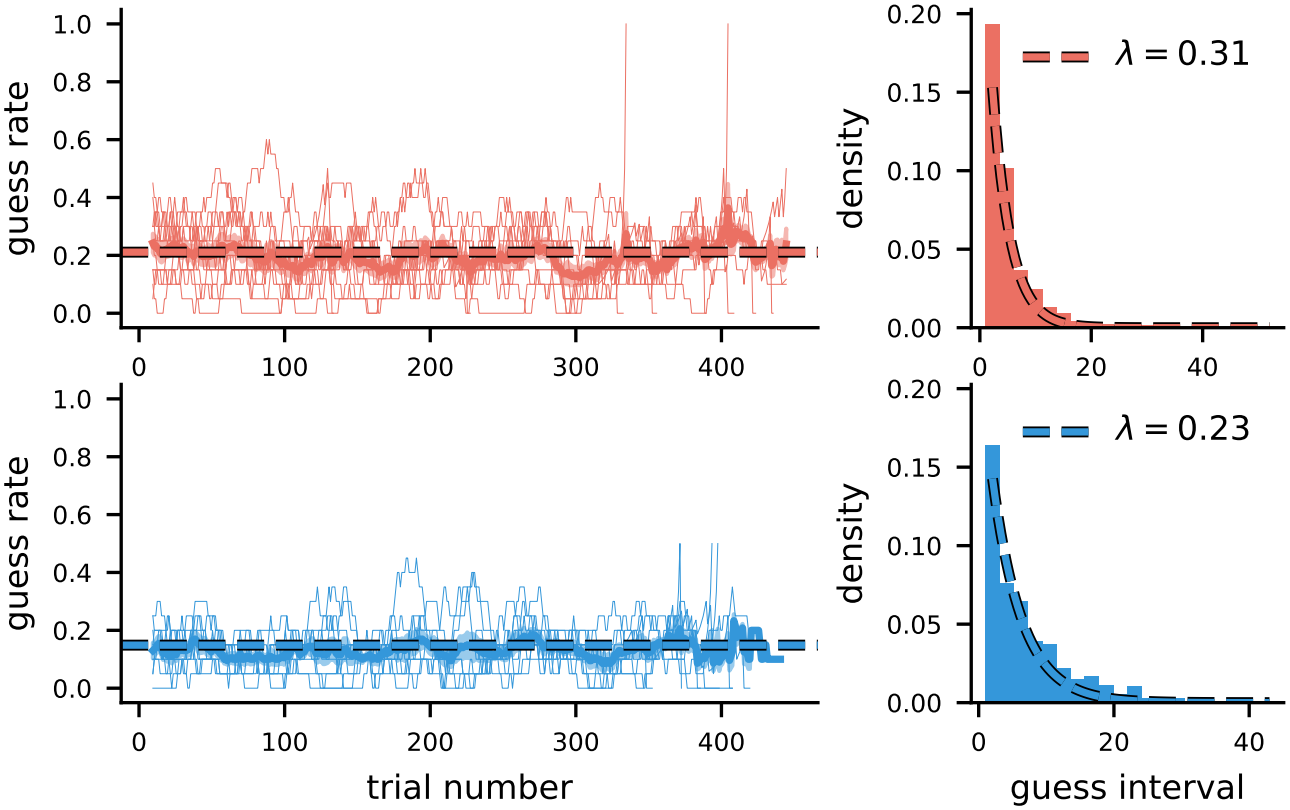
The dynamics of the guess rate across sessions. The rate of guess trials (left) and inter-guess-interval (right) across sessions for monkey E (top) and W (bottom).

Each group can then be described by its preferred stimulus, *x*. One of these groups, the local group, includes the target stimulus, and the rest do not; the goal is to compute the chances one of these non local groups has a higher firing rate than the local one. Assuming Gaussian noise in both the non-local and local groups, the probability of a group *i* beats the local group is

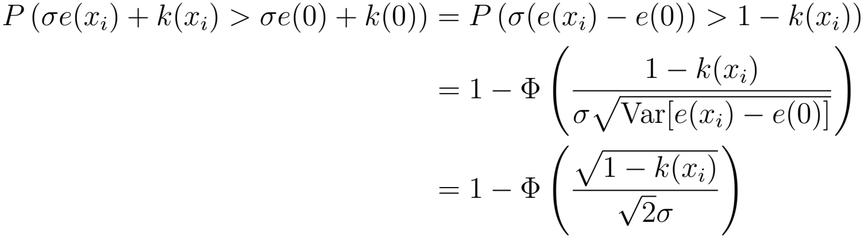

where Φ is the standard Gaussian CDF. The probability of the union of all these events is intractable to compute, but can be upper bounded by ignoring the intersections and just taking a sum: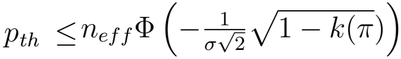, since, by construction, *k*(*x*_*i*_) *≈ k*(*π*) for all the non-local groups. When each event is sufficiently rare, this upper bound is a fine approximation for *p*_*th*_.

But the union bound becomes too loose in the presence of high noise and severely overestimates the number of threshold errors. Practically speaking, it exceeds 1 and is no longer a valid probability. More fundamentally, the probability of each non-local region beating the local one becomes too large to be treated as disjoint events. Instead, we can compute the probability of no off-target group winning, the joint probability *P* (*e*(*x*) *− e*(0) *>* 1 *− k*(*x*) *∀x*). The noise differences, *e*(*x*) *− e*(0), are jointly Gaussian and correlated, and so this probability is also intractable in general. While the differences are correlated, we can approximate them as conditionally independent given the noise of the local group: _say that 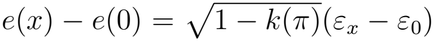_ for independent standard normals, *ε*. The choice of scale, 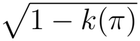, ensures that this approximation matches the marginal variances. In that case the probability is 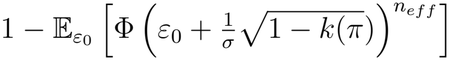. The expectation requires a numerical integration over *ε*_0_, but this is at least an improvement from the naive calculation.

Finally, there is a subtle difference between ‘threshold errors’, as discussed here, and the guess rate of a mixture model. Namely, guesses in the mixture model produce a uniform distribution over the circle, while threshold errors produce a uniform distribution only among the non-local groups. To convert from one to the other, we fix the local component, *p*_*c*_, to have an MCE matching the FI-derived upper bound, and then find the guess rate, *p*_*g*_, such that the mixture’s density at 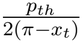. With that in hand, the total error predicted by all these approximations is

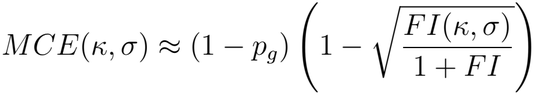

**Figure S4:**
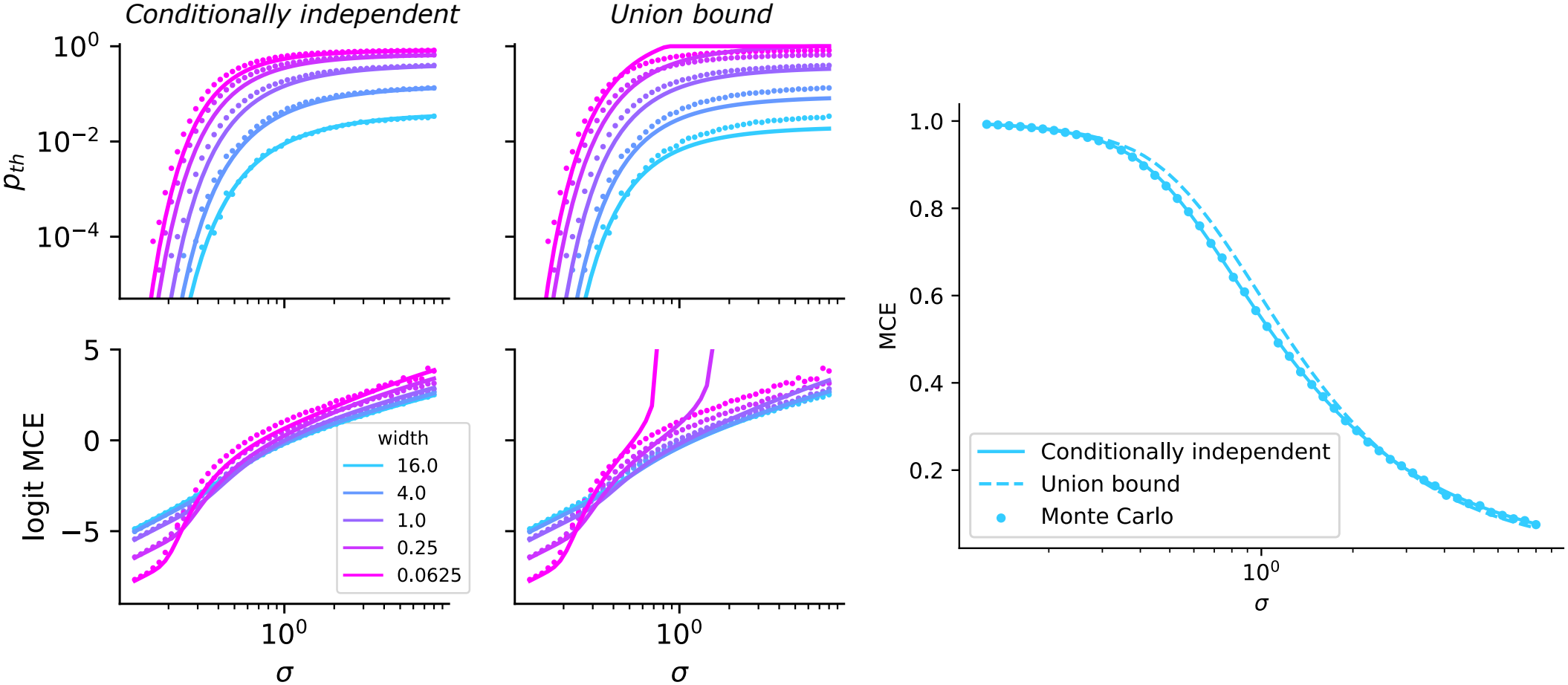
Comparison of approximation schemes. On the left is our approximation based on the expected product of conditional probabilities. In the middle, is the modified union bound approximation, based on the sum of marginal probabilities. The top row shows *p*_*th*_ predicted by each approximation, as well as that computed empirically via monte carlo (showing the proportion of errors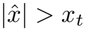). On the bottom, is the MCE plotted as the logit (i.e. log(*x*)*−*log(1*−x*)) to help see the extremes on either end. On the right, we show the MCE just for the wide (nearly linear) kernel, to remove the crowding of other lines.

**Figure S5:**
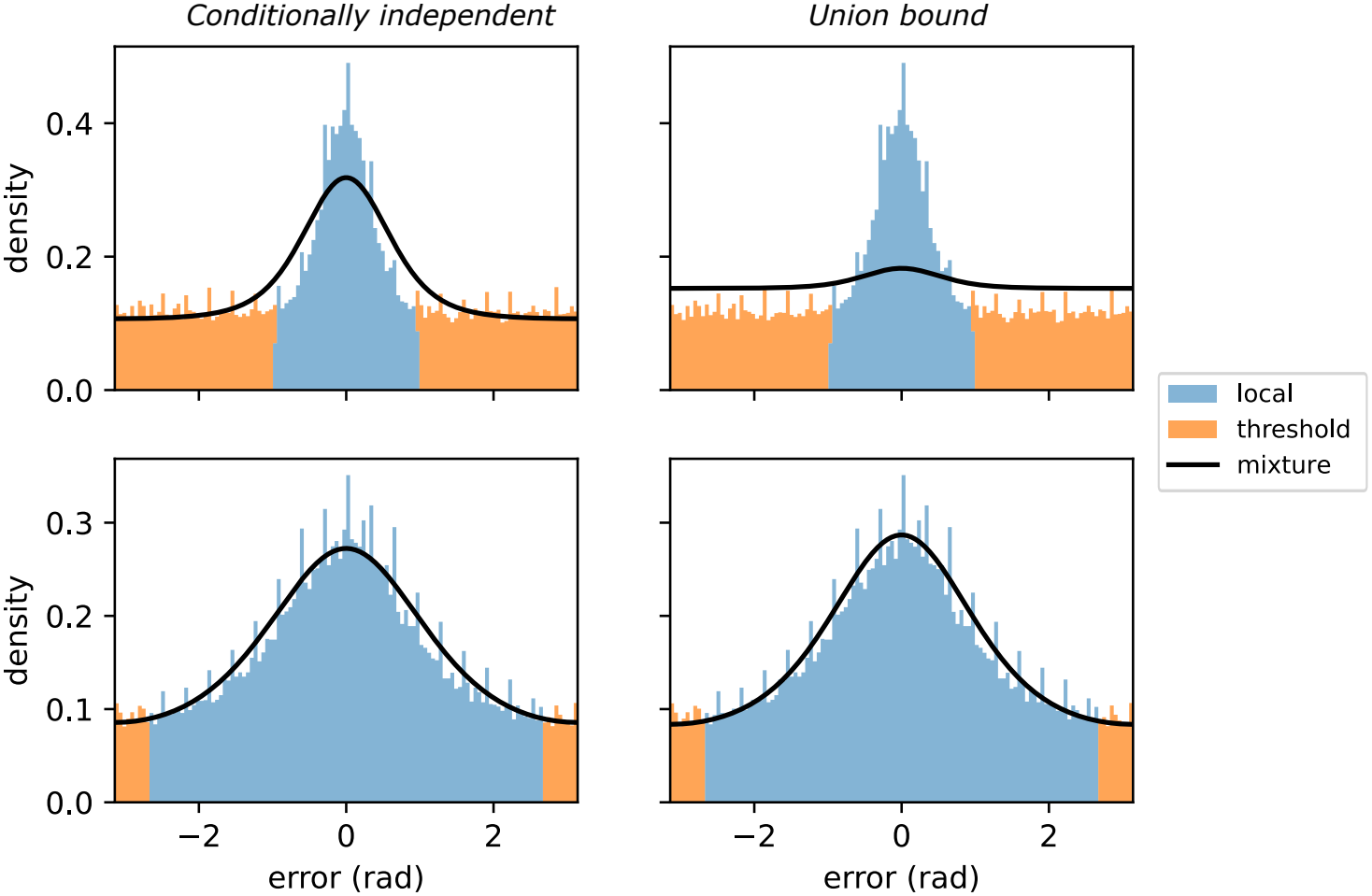
Examples of the 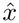 distributions for two different kernel widths (top: *κ* = 4, bottom: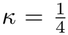) and one noise level (*σ* = 2), as well as the predicted mixture model from each approximation scheme. The area that is considered a ‘threshold’ error, i.e.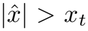, is shown in orange. That is, *p*_*th*_ computes the mass of the orange area. The histograms come from from 50 thousand noise samples evaluated on 500 color bins.

**Figure S6:**
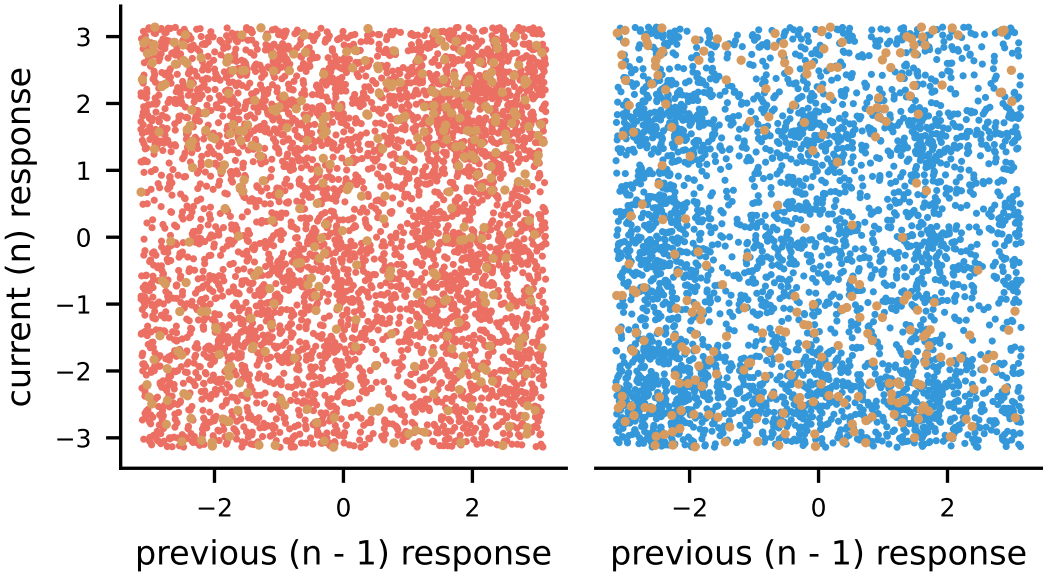
The correlation of responses on subsequent trials. The previous (or trial n - 1) versus current (or trial n) responses for monkey E (left) and W (right).

**Figure S7:**
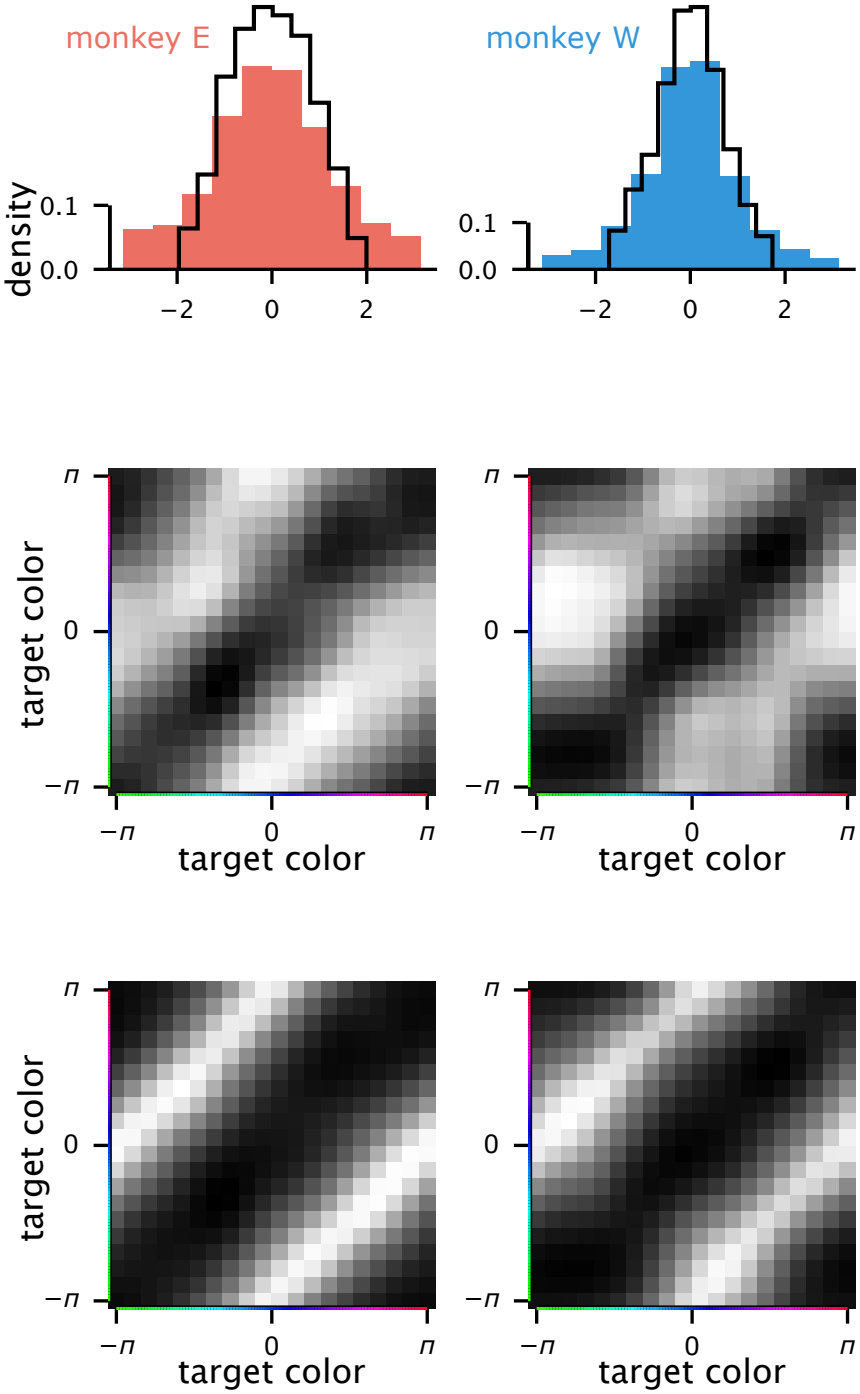
(top) The histogram of behavioral errors for both the full data (color) and data selected as correct for the geometry estimations (black line). (middle) The smoothed response map for target-response color pairs for the full data and (bottom) the trials selected as correct.

**Figure S8:**
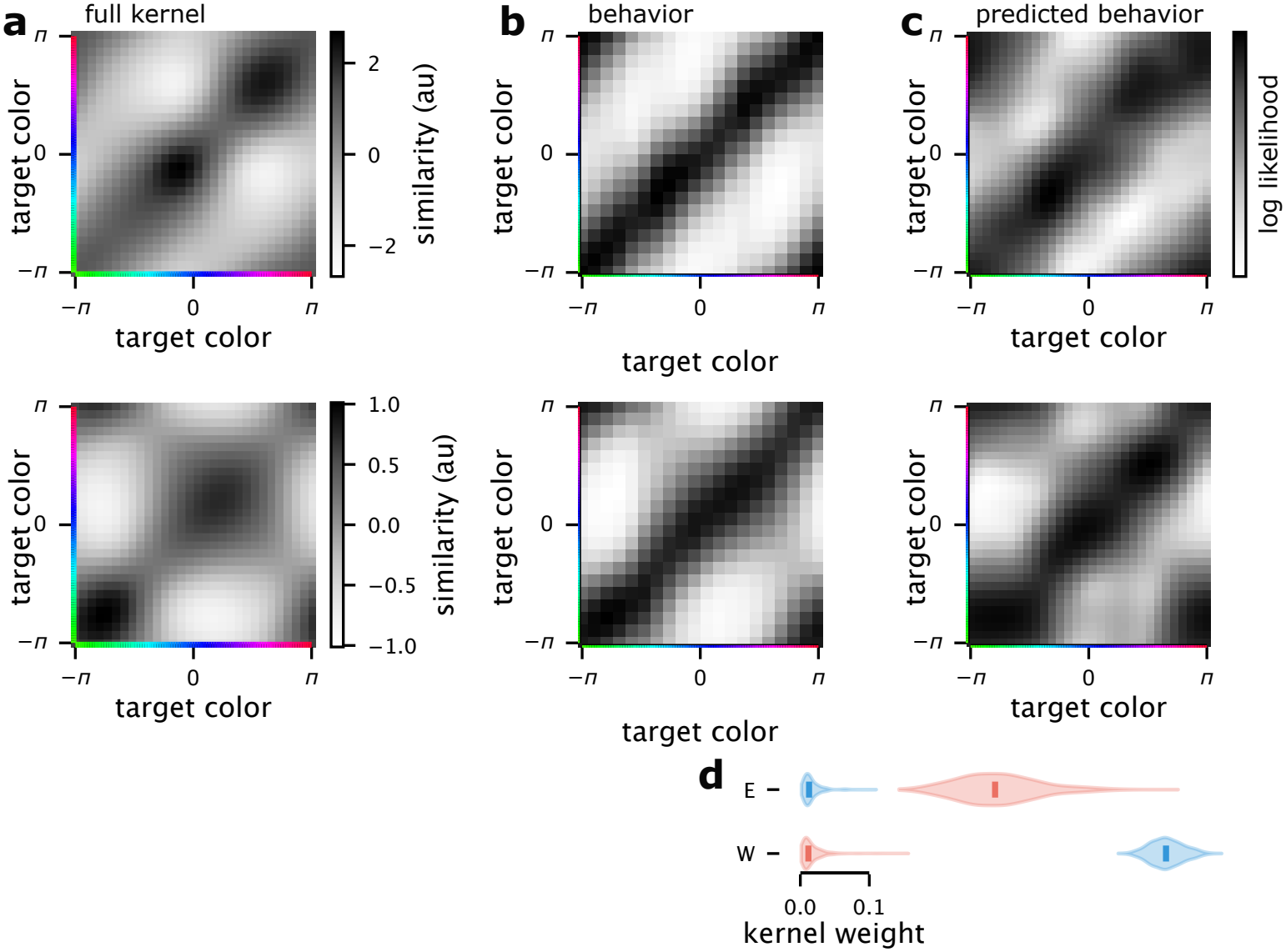
Results for the full kernel with trials resampled to have identical response-target distributions. a - d are the same as fig. 5f - i, but for the restricted trial set. The resampling was performed by discretizing the response deviations into 10 bins, taking only the bins with error less than *π/*4 and then randomly undersampling (as implemented in [52]) so that every bin has the same number of trials.

## References

1. Van Den Berg, R., Shin, H., Chou, W.-C., George, R. & Ma, W. J. Variability in Encoding Precision Accounts for Visual Short-Term Memory Limitations. Proceedings of the National Academy of Sciences 109, 8780–8785. issn: 0027-8424, 1091-6490. (2024) (May 2012).

2. McMaster, J. M., Tomić, I., Schneegans, S. & Bays, P. M. Swap Errors in Visual Working Memory Are Fully Explained by Cue-Feature Variability. Cognitive Psychology 137, 101493. issn: 00100285. (2024) (Sept. 2022).

3. Van Den Berg, R., Awh, E. & Ma, W. J. Factorial Comparison of Working Memory Models. Psychological Review 121, 124–149. issn: 1939-1471, 0033-295X. (2024) (2014).

4. Rajsic, J., Swan, G., Wilson, D. E. & Pratt, J. Accessibility Limits Recall from Visual Working Memory. Journal of Experimental Psychology: Learning, Memory, and Cognition 43, 1415–1431. issn: 1939-1285, 0278-7393. (2024) (Sept. 2017).

5. Bays, P., Schneegans, S., Ma, W. J. & Brady, T. F. Representation and Computation in Working Memory Oct. 2022. (2025).

6. Bays, P. M., Catalao, R. F. G. & Husain, M. The Precision of Visual Working Memory Is Set by Allocation of a Shared Resource. Journal of Vision 9, 7–7. issn: 1534-7362. (2024) (Sept. 2009).

7. Zhang, W. & Luck, S. J. Discrete Fixed-Resolution Representations in Visual Working Memory. Nature 453, 233–235. issn: 0028-0836, 1476-4687. (2024) (May 2008).

8. Schurgin, M. W., Wixted, J. T. & Brady, T. F. Psychophysical Scaling Reveals a Unified Theory of Visual Memory Strength. Nature Human Behaviour 4, 1156–1172. issn: 2397-3374. (2024) (Sept. 2020).

9. Ma, W. J., Husain, M. & Bays, P. M. Changing Concepts of Working Memory. Nature Neuroscience 17, 347–356. issn: 1097-6256, 1546-1726. (2026) (Mar. 2014).

10. Alleman, M., Panichello, M., Buschman, T. J. & Johnston, W. J. The Neural Basis of Swap Errors in Working Memory. Proceedings of the National Academy of Sciences 121, e2401032121. issn: 0027-8424, 1091-6490. (2025) (Aug. 2024).

11. Tomić, I. & Bays, P. M. Perceptual Similarity Judgments Do Not Predict the Distribution of Errors in Working Memory. Journal of Experimental Psychology: Learning, Memory, and Cognition 50, 535–549. issn: 1939-1285, 0278-7393. (2025) (Apr. 2024).

12. Wimmer, K., Nykamp, D. Q., Constantinidis, C. & Compte, A. Bump Attractor Dynamics in Prefrontal Cortex Explains Behavioral Precision in Spatial Working Memory. Nature Neuroscience 17, 431–439. issn: 1097-6256, 1546-1726. (2026) (Mar. 2014).

13. Panichello, M. F. & Buschman, T. J. Shared Mechanisms Underlie the Control of Working Memory and Attention. Nature 592, 601–605. issn: 0028-0836, 1476-4687. (2024) (Apr. 2021).

14. Adam, K. C., Vogel, E. K. & Awh, E. Clear Evidence for Item Limits in Visual Working Memory. Cognitive Psychology 97, 79–97. issn: 00100285. (2026) (Sept. 2017).

15. Curtis, C. E. & D’Esposito, M. Persistent Activity in the Prefrontal Cortex during Working Memory. Trends in Cognitive Sciences 7, 415–423. issn: 13646613. (2026) (Sept. 2003).

16. Constantinidis, C. et al. Persistent Spiking Activity Underlies Working Memory. The Journal of Neuroscience 38, 7020–7028. issn: 0270-6474, 1529-2401. (2026) (Aug. 2018).

17. Panichello, M. F. et al. Intermittent Rate Coding and Cue-Specific Ensembles Support Working Memory. Nature. issn: 0028-0836, 1476-4687. (2024) (Nov. 2024).

18. Lundqvist, M., Herman, P. & Miller, E. K. Working Memory: Delay Activity, Yes! Persistent Activity? Maybe Not. The Journal of Neuroscience 38, 7013–7019. issn: 0270-6474, 1529-2401. (2026) (Aug. 2018).

19. Barbosa, J. et al. Interplay between Persistent Activity and Activity-Silent Dynamics in the Pre-frontal Cortex Underlies Serial Biases in Working Memory. Nature Neuroscience 23, 1016–1024. issn: 1097-6256, 1546-1726. (2026) (Aug. 2020).

20. Xie, Y. et al. Geometry of Sequence Working Memory in Macaque Prefrontal Cortex. Science 375, 632–639. issn: 0036-8075, 1095-9203. (2024) (Feb. 2022).

21. Buschman, T. J., Siegel, M., Roy, J. E. & Miller, E. K. Neural Substrates of Cognitive Capacity Limitations. Proceedings of the National Academy of Sciences 108, 11252–11255. issn: 0027-8424, 1091-6490. (2026) (July 2011).

22. Wu, S., Hamaguchi, K. & Amari, S.-i. Dynamics and Computation of Continuous Attractors. Neural Computation 20, 994–1025. issn: 0899-7667, 1530-888X. (2026) (Apr. 2008).

23. Fung, C. C. A., Wong, K. Y. M. & Wu, S. A Moving Bump in a Continuous Manifold: A Comprehensive Study of the Tracking Dynamics of Continuous Attractor Neural Networks. Neural Computation 22, 752–792. issn: 0899-7667, 1530-888X. (2026) (Mar. 2010).

24. Burak, Y. & Fiete, I. R. Fundamental Limits on Persistent Activity in Networks of Noisy Neurons. Proceedings of the National Academy of Sciences 109, 17645–17650. issn: 0027-8424, 1091-6490. (2024) (Oct. 2012).

25. Wei, X.-X. & Woodford, M. Representational Geometry Explains Puzzling Error Distributions in Behavioral Tasks Jan. 2023. (2024).

26. Kim, J. H. J., Fiete, I. & Schwab, D. J. Superlinear Precision and Memory in Simple Population Codes Aug. 2020. arXiv: 2008.00629 [q-bio]. (2024).

27. Bays, P. M. Noise in Neural Populations Accounts for Errors in Working Memory. The Journal of Neuroscience 34, 3632–3645. issn: 0270-6474, 1529-2401. (2024) (Mar. 2014).

28. Blanco Malerba, S., Pieropan, M., Burak, Y. & Azeredo Da Silveira, R. Random Compressed Coding with Neurons. Cell Reports 44, 115412. issn: 22111247. (2026) (Mar. 2025).

29. Johnston, W. J. & Freedman, D. J. Redundant Representations Are Required to Disambiguate Simultaneously Presented Complex Stimuli. PLOS Computational Biology 19 (ed Yang, T.) e1011327. issn: 1553-7358. (2026) (Aug. 2023).

30. Stringer, C., Michaelos, M., Tsyboulski, D., Lindo, S. E. & Pachitariu, M. High-Precision Coding in Visual Cortex. Cell 184, 2767–2778.e15. issn: 00928674. (2024) (May 2021).

31. Noorman, M., Hulse, B. K., Jayaraman, V., Romani, S. & Hermundstad, A. M. Maintaining and Updating Accurate Internal Representations of Continuous Variables with a Handful of Neurons. Nature Neuroscience 27, 2207–2217. issn: 1097-6256, 1546-1726. (2026) (Nov. 2024).

32. Bays, P. M. Spikes Not Slots: Noise in Neural Populations Limits Working Memory. Trends in Cognitive Sciences 19, 431–438. issn: 13646613. (2026) (Aug. 2015).

33. Schneegans, S. & Bays, P. M. Neural Architecture for Feature Binding in Visual Working Memory. The Journal of Neuroscience 37, 3913–3925. issn: 0270-6474, 1529-2401. (2024) (Apr. 2017).

34. Luck, S. J. & Vogel, E. K. Visual Working Memory Capacity: From Psychophysics and Neurobiology to Individual Differences. Trends in Cognitive Sciences 17, 391–400. issn: 13646613. (2026) (Aug. 2013).

35. Luck, S. J. & Vogel, E. K. The Capacity of Visual Working Memory for Features and Conjunctions. Nature 390, 279–281. issn: 0028-0836, 1476-4687. (2026) (Nov. 1997).

36. Noorman, M., Hulse, B. K., Jayaraman, V., Romani, S. & Hermundstad, A. M. Accurate Angular Integration with Only a Handful of Neurons Preprint (Neuroscience, May 2022). (2024).

37. Ben-Yishai, R., Bar-Or, R. L. & Sompolinsky, H. Theory of Orientation Tuning in Visual Cortex. Proceedings of the National Academy of Sciences 92, 3844–3848. issn: 0027-8424, 1091-6490. (2024) (Apr. 1995).

38. Skaggs, W. E., Knierim, J. J., Kudrimoti, H. S. & McNaughton, B. L. A Model of the Neural Basis of the Rat’s Sense of Direction.

39. Zhang, K. Representation of Spatial Orientation by the Intrinsic Dynamics of the Head-Direction Cell Ensemble: A Theory. The Journal of Neuroscience 16, 2112–2126. issn: 0270-6474, 1529-2401. (2026) (Mar. 1996).

40. Deng, Z., Shi, J. & Zhu, J. NeuralEF: Deconstructing Kernels by Deep Neural Networks Oct. 2022. arXiv: 2205.00165 [cs]. (2026).

41. Courellis, H. S. et al. Abstract Representations Emerge in Human Hippocampal Neurons during Inference. Nature 632, 841–849. issn: 0028-0836, 1476-4687. (2025) (Aug. 2024).

42. Bernardi, S. et al. The Geometry of Abstraction in the Hippocampus and Prefrontal Cortex. Cell 183, 954–967.e21. issn: 00928674. (2024) (Nov. 2020).

43. Nogueira, R., Rodgers, C. C., Bruno, R. M. & Fusi, S. The Non-Linear Mixed Representations in Somatosensory Cortex Support Simple and Complex Tasks.

44. Rigotti, M. et al. The Importance of Mixed Selectivity in Complex Cognitive Tasks. Nature 497, 585–590. issn: 0028-0836, 1476-4687. (2024) (May 2013).

45. Johnston, W. J. & Fusi, S. Abstract Representations Emerge Naturally in Neural Networks Trained to Perform Multiple Tasks. Nature Communications 14, 1040. issn: 2041-1723. (2024) (Feb. 2023).

46. Panichello, M. F., DePasquale, B., Pillow, J. W. & Buschman, T. J. Error-Correcting Dynamics in Visual Working Memory. Nature Communications 10, 3366. issn: 2041-1723. (2024) (July 2019).

47. Wang, Z., Stocker, A. A. & Lee, D. D. Efficient Neural Codes That Minimize L p Reconstruction Error. Neural Computation 28, 2656–2686. issn: 0899-7667, 1530-888X. (2024) (Dec. 2016).

48. Wang, Z., Wei, X.-X., Stocker, A. A. & Lee, D. D. Efficient Neural Codes under Metabolic Constraints.

49. Zhang, K. & Sejnowski, T. J. Neuronal Tuning: To Sharpen or Broaden? Neural Computation 11, 75–84. issn: 0899-7667, 1530-888X. (2024) (Jan. 1999).

50. Pedregosa, F. et al. Scikit-Learn: Machine Learning in Python. MACHINE LEARNING IN PYTHON.

51. Mardia, K. V. & Jupp, P. E. Directional Statistics isbn: 978-0-471-95333-3 (J. Wiley, Chichester; New York, 2000).

52. Lemaıtre, G. & Nogueira, F. Imbalanced-Learn: A Python Toolbox to Tackle the Curse of Imbalanced Datasets in Machine Learning.

